# Epistatic networks jointly influence phenotypes related to metabolic disease and gene expression in Diversity Outbred mice

**DOI:** 10.1101/098681

**Authors:** Anna L. Tyler, Bo Ji, Daniel M. Gatti, Steven C. Munger, Gary A. Churchill, Karen L. Svenson, Gregory W. Carter

## Abstract

Genetic studies of multidimensional phenotypes can potentially link genetic variation, gene expression, and physiological data to create multi-scale models of complex traits. Multi-parent populations provide a resource for developing methods to understand these relationships. In this study, we simultaneously modeled body composition, serum biomarkers, and liver transcript abundances from 474 Diversity Outbred mice. This population contained both sexes and two dietary cohorts. Using weighted gene co-expression network analysis (WGCNA), we summarized transcript data into functional modules which we then used as summary phenotypes representing enriched biological processes. These module phenotypes were jointly analyzed with body composition and serum biomarkers in a combined analysis of pleiotropy and epistasis (CAPE), which inferred networks of epistatic interactions between quantitative trait loci that affect one or more traits. This network frequently mapped interactions between alleles of different ancestries, providing evidence of both genetic synergy and redundancy between haplotypes. Furthermore, a number of loci interacted with sex and diet to yield sex-specific genetic effects. We were also able to identify alleles that potentially protect individuals from the effects of a high-fat diet. Although the epistatic interactions explained small amounts of trait variance, the combination of directional interactions, allelic specificity, and high genomic resolution provided context to generate hypotheses for the roles of specific genes in complex traits. Our approach moves beyond the cataloging of single loci to infer genetic networks that map genetic etiology by simultaneously modeling all phenotypes.

---

Deriving biological models from genetic studies with multi-dimensional phenotype data requires analytical methods that distill the complexity of genetic systems to specific hypotheses. This challenge has become increasingly acute with the advent of genome-scale data resources designed to determine how genetic variation affects biological processes at molecular resolution. Genetic studies of gene expression (Hemani *et al.* 2014; Schadt *et al.* 2003; Chesler *et al.* 2005), protein expression (Picotti *et al.* 2013; Chick *et al.* 2016), and other panels of quantitative traits (Jia and Jannink 2012; Wolf *et al.* 2006) can potentially map the path from genetic variants to complex physiological states through dysregulated processes and pathways (Civelek *et al.* 2017; Albert and Kruglyak 2015). Traits related to metabolic disease, such as obesity and blood lipid profiles, are examples of such a system (Schork 1997). Many genetic factors, which possibly interact, influence multiple traits including molecular regulation, serum composition, and health outcomes. Identifying these genes and their interactions will play a critical role in predicting individual susceptibility to metabolic disease and prioritizing drug targets for targeted treatments (Moore and Williams 2009). However, despite availability of large-scale studies in multiple human populations, little is known about the genetic architecture of metabolic disease. This is likely due to a number of factors, including variable environmental exposure and structured populations (Rosenberg *et al.* 2002) that affect allele frequencies (Greene *et al.* 2009; Pritchard *et al.* 2000). The analysis of epistatic networks is further complicated by the fact that epistasis contributes to additive variance (Huang and Mackay 2016).

Multi-parent populations of model organisms, such as the Diversity Outbred (DO) mice (Svenson *et al.* 2012), offer a powerful alternative for mapping the genetic architecture of complex traits. This outbred population contains extensive allelic variation drawn from both classic and more recently wild-derived inbred strains, which is evenly distributed across the genome (Philip *et al.* 2014; Svenson *et al.* 2012; Logan *et al.* 2013). The resulting density of polymorphisms enables a much higher resolution of mapping than typical intercross populations that share large regions of common ancestry (Yang *et al.* 2011). Furthermore, the breeding paradigm in the DO is designed to maintain allelic diversity, reduce linkage disequilibrium, and generate minimal population structure (Svenson *et al.* 2012; Chesler *et al.* 2016). The resulting allelic balance allows mapping to narrow genomic loci and can potentially power studies of epistasis. Indeed, although many heritable phenotypes have been measured in DO mice (Svenson *et al.* 2012; Bogue *et al.* 2015), these studies rarely identify a single QTL of exceptional effect (Churchill *et al.* 2012; Logan *et al.* 2013). The DO is therefore amenable to genetic analysis but often too complex for standard single-trait QTL mapping.

The proliferation of expression QTL (eQTL) and similar studies to link genome-wide molecular traits with genetic variation has generated new strategies for trait mapping. A key advance has been the mapping of dimensionally-reduced representations of these data to yield concise associations between QTL and summary phenotypes. This strategy has been used to prioritize potential regulator genes based on genetic association and co-expressed genes (Biswas *et al.* 2008; Ghazalpour *et al.* 2006), protein expression (Chick *et al.* 2016), and correlated phenotypes (Neto *et al.* 2008). The precision of these molecular traits has also powered the detection of epistasis, including in human studies (Lappalainen *et al.* 2011; Hemani *et al.* 2014). In total, these studies have magnified the power of genetic studies to identify the pathways and processes that underlie organism-level phenotypes. This study advances this integrative analysis strategy by inferring epistatic networks of interacting QTL that selectively influence multiple physiological, serum, and gene module traits. These QTL, derived from haplotype association, map how alleles from each inbred founder strain interact with each other, sex, and diet to affect these multiple related traits. In contrast to similar analyses of intercross data (Tyler *et al.* 2016), distinctions can be drawn between inter-strain and intra-strain epistasis, which suggested instances of both cross-strain incompatibilities and within-strain synergy. We used the network structure to identify candidate genes and derive hypotheses of complex trait regulation. Our approach, based on the combined analysis of pleiotropy and epistasis (CAPE) (Tyler *et al.* 2013), is generalizable to all multi-parent populations with high dimensional phenotype data.

## Materials and Methods

The primary goal of this study was to determine how multiple QTL interact to affect complex traits in DO mice. To do this, we followed a work flow with four major phases (Figure 1). First, we collected data on multiple metabolic traits from mice fed with chow or high fat diets. Second, physiological traits and RNA-seq data were batch corrected and further processed to produce the inputs used for epistasis inference. Third, we developed and applied a multi-parent version of our previously described software for combined analysis of pleiotropy and epistasis (CAPE). This analysis pipeline combines information across multiple traits to infer directed epistasis between genetic variants. Fourth, we further interpreted the epistatic interactions that formed a connected network between founder haplotypes. We assessed intra- and inter-strain interactions, considered QTL interactions with sex and diet, and analyzed the overall network structure.

**Figure 1.**
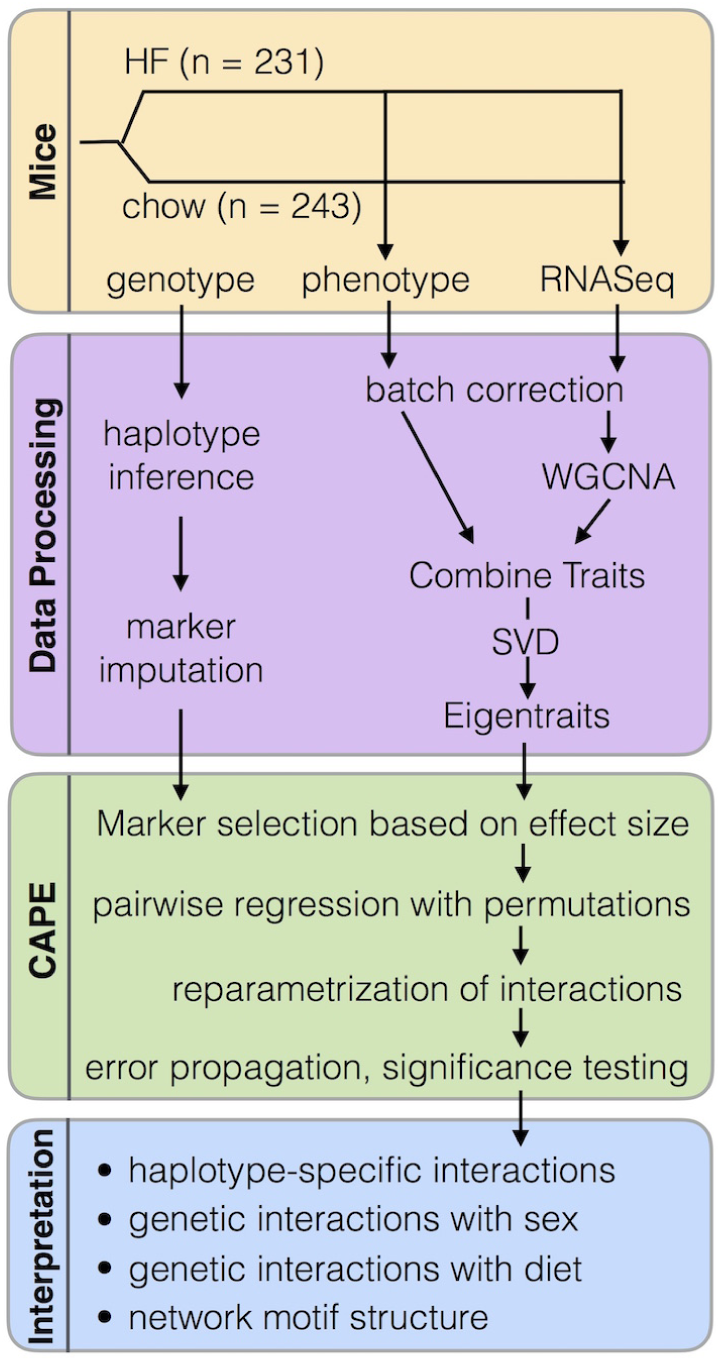
Overview of study work flow. Boxes separate steps into four themes: mouse experimentation, processing of genotype and phenotype data, combined analysis of pleiotropy and epistasis (CAPE), and interpretation of network results.

### Mice

Mice were obtained from The Jackson Laboratory (Bar Harbor, ME) as described in (Svenson *et al.* 2012). The animals were non-sibling DO mice ranging from generation 4 to 11, and males and females were represented equally. All animal procedures were approved by the Animal Care and Use Committee at The Jackson Laboratory (Animal Use Summary # 06006). Mice were housed in same-sex cages with five animals per cage as described in (Svenson *et al.* 2012). Animals had free access to either standard rodent chow (6% fat by weight, LabDiet 5K52, LabDiet, Scott Distributing, Hudson, NH) or a high-fat, high-sucrose (HF) diet (Envigo Teklad TD.08811, Envigo, Madison, WI) for the duration of the study protocol (26 weeks). Caloric content of the HF diet was 45% fat, 40% carbohydrates and 15% protein. Diets were assigned randomly.

#### Genotyping

Genotyping was performed on tail biopsies as described in (Svenson *et al.* 2012) using the Mouse Universal Genotyping Array (MUGA) (7,854 markers) and the MegaMUGA (77,642 markers) (GeneSeek, Lincoln, NE).

#### Measurement of physiological traits

Physiological traits were measured as described in (Svenson *et al.* 2012). Blood was collected retro-orbitally in 10-week old mice after administration of local anesthetic. Cholesterol and triglycerides were measured using the Beckman Synchron DXC600Pro Clinical chemistry analyzer. Leptin was measured in 8-week old mice using non-fasted plasma prepared as previously described (Svenson *et al.* 2012). Leptin levels were analyzed using the Meso Scale Discovery electrochemiluminescent system according to the manufacturer’s recommended protocol (Meso Scale Diagnostics, Rockville, MD). Body composition (lean mass and total mass) were measured in 12-week old mice by dual X-ray absorptiometry (DEXA) using a Lunar PIXImus densitometer (GE Medical Systems). Fat mass was calculated as log(total mass - lean mass).

#### Measurement of transcript abundance

We measured transcriptome-wide expression levels from whole livers as described in (Chick *et al.* 2016; Munger *et al.* 2014). We sequenced RNA using single-end RNA-Seq (Munger *et al.* 2014), and aligned transcripts to strain-specific genomes from the DO founders (Chick *et al.* 2016). We used an expectation maximization algorithm (EMASE, https://github.com/churchill-lab/emase) to estimate read counts. We corrected for the effects of sex, diet, and batch by normalizing read counts in each sample using upper-quantile normalization and applying a rank Z transformation across samples.

### Data Processing

#### Founder haplotype inference

The MUGA and MegaMUGA arrays identify single nucleotide polymorphisms (SNPs) present in each individual. We converted the SNP calls from the arrays to founder haplotypes. We did this using a hidden Markov model (HMM) (Gatti *et al.* 2014) which uses the order of SNPs in an individual mouse to infer transition points between different DO founder haplotypes. The result is a probability of each parental haplotype at each SNP position in the genome (Gatti *et al.* 2014). We also merged diplotype probabilities from the MUGA and MegaMUGA to interpolate markers on an evenly spaced 64,000 marker grid (0.0238 cM between markers). A few samples were unable to be genotyped using either the MUGA or MegaMUGA. Theses samples were genotyped using genotyping by RNA-sequence (GBRS) (Chick *et al.* 2016). GBRS reconstructs individual genotypes from RNA-Seq data without using genotyping arrays. The method aligns RNA-Seq reads to a common pooled transcriptome of all founder strains simultaneously and matches the array calls with high fidelity. The mean Pearson correlation between GBRS genotypes and array genotypes is 0.88 (standard deviation 0.03). The software package is freely available at https://github.com/churchill-lab/gbrs.

#### Transcript filtering

Because we were interested in epistatic interactions influencing transcription, we filtered the liver transcripts to a subset that were likely to be influenced by multiple loci (Figure 2). To do this, we identified transcripts that were likely to have both a local (*cis*) and a distant (*trans*) expression QTL (eQTL). First, we used DOQTL (Gatti *et al.* 2014) to identify *cis*-eQTL for all transcripts that were expressed in at least 50 animals (26,875 transcripts). We corrected for sex, diet, and batch and used hierarchical linear models to correct for genetic relatedness (Kang *et al.* 2008). We kept all transcripts (13,228) with a significant *cis*-eQTL, which we defined as a significant eQTL (permutation-based *p* ≤ 0.05, LOD ≥ 7.4) within 2 Mbp of the encoding gene’s transcript. To identify transcripts with *trans* effects, we regressed out the effects of the *cis*-eQTL for each transcript (Pierce *et al.* 2014) and re-mapped QTL using DOQTL. We kept all (3,635) transcripts with a significant eQTL (permutation-based p value *≤* 0.05, LOD *>*= 7.4) at least 10 Mb away from the encoding gene. To test the robustness of our filtering methods, we re-ran this pipeline using p value thresholds of 0.01 (1719 final transcripts), 0.1 (6253 final transcripts), and 0.2 (10,000 final transcripts). As we describe below, different filtering thresholds yielded similar results.

**Figure 2.**
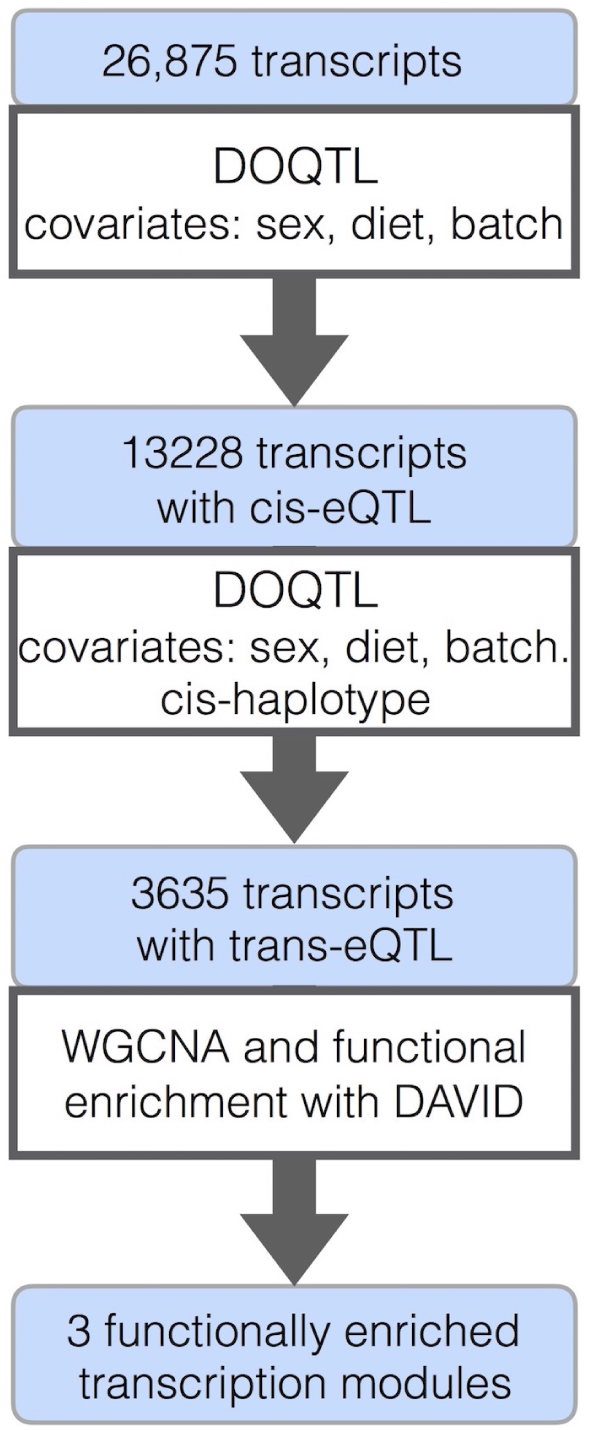
Overview of methods used to filter transcripts with potential *trans*-eQTL and create co-expression modules.

#### Transcript clustering

After identifying transcripts with likely *trans*-eQTL, we collapsed the 3,635 transcripts into summary expression traits using Weighted Gene Co-expression Network Analysis (WGCNA) (Langfelder and Horvath 2008, 2012; R Core Team 2016). This step removes the burden of interpreting epistatic interactions influencing thousands of individual transcripts by condensing the transcripts into functionally enriched modules representing transcriptional programs (Zhao *et al.* 2010; Fuller *et al.* 2007), for example immune processes or redox reactions. Whereas other transcript clustering methods, such as singular value decomposition (SVD) (Alter *et al.* 2000) and independent component analysis (ICA) (Liebermeister 2002; Rotival *et al.* 2011) group genes purely on the statistical properties of expression matrices, WGCNA treats correlations between transcripts as connections in a network, which has been shown to be biologically relevant (Barabási and Oltvai 2004; Featherstone and Broadie 2002; Agrawal 2002).

WGCNA was used to cluster hepatic transcripts with likely *trans* effects into functional modules. We ran WGCNA on the 3,635 transcripts with likely *trans*-eQTL using default settings. The analysis yielded 11 gene modules, three of which had significantly enriched functions: metabolic processes, redox reactions, and immune processes (*p* ≤ 0.05, with Benjamini correction for multiple comparisons) (Huang *et al.* 2009a,b) (Table S1). We refer to the modules by their functional enrichments: the metabolism module, the redox module, and the immune module. All three modules were recapitulated by WGCNA at significance thresholds of *p* ≤ 0.1 and *p* ≤ 0.2, and both the immune and metabolism modules were recapitulated for *p* ≤ 0.01, indicating that the transcript clustering is robust to different thresholds used during transcript filtering. The Pearson correlation (*r*) between the original expression modules and the expression modules at different thresholds ranged from *r* = 0.85 to *r* = 0.95. The functional enrichments found here were similar to those found in previous work using WGCNA to analyze mouse hepatic transcripts (Liu and Ye 2014).

Following standard practice, we used the module *eigengene* (first principal component) to represent the three enriched modules in our analysis. Although these expression traits do not represent individual transcripts, they represent transcriptional programs that are potentially relevant to the physiological phenotypes we analyzed here.

#### Combining expression and physiological traits

We combined the transcription modules from WGCNA with our physiological traits in order to identify genetic interactions influencing both transcriptional programs and physiological traits simultaneously. We rank Z normalized each of the physiological traits. Fat mass was log-transformed to reproduce a more linear relationship with lean mass (Forbes 1987). We then combined the physiological traits with the transcript module eigengenes from WGCNA and performed singular value decomposition (SVD) on the trait matrix to obtain eight orthogonal eigentraits (ETs). The ETs combine common signals across all traits and may group functionally related signals into individual ETs. This concentration of functional effects may improve power to map weak effects that are distributed across multiple traits. In our analysis effect sizes of markers influencing ETs were comparable to those of the individual traits, but identified different significant QTL (Results).

### Combined analysis of pleiotropy and epistasis

We previously developed CAPE to infer directed genetic interactions by combining information across multiple phenotypes (Carter *et al.* 2012; Tyler *et al.* 2013). For the analysis in the DO, we adapted CAPE to infer epistasis in multiparental populations. With these changes CAPE can be applied to DO mice as well as other multiparental populations, including in other model organisms. It can also be used to analyze interactions between single nucleotide polymorphisms (SNPs) in human populations. To adapt CAPE to multiparental populations we made two major changes. First, we use an (n-1)-state model to estimate individual haplotype effects at each locus. In the DO there are eight possible haplotypes, derived from the eight DO founders, at each locus. We used a seven-state linear model to estimate the effects of each allele using the B6 haplotype as the reference haplotype. Thus all effects are in reference to B6, and B6 does not explicitly appear in any QTL or the epistatic network. The second major change we made to CAPE was to specify epistatic interactions in terms of ancestral haplotype. We report, for example, and interaction between the A/J haplotype on Chr 9 and the CAST haplotype on Chr 2, rather than simply an interaction between unlabeled alternate alleles on Chrs 9 and 2.

For phenotypes, we chose to use the first three ETs. We considered a range of two to six ETs and empirically determined that power to detect interactions is roughly equal for the first three, four, or five ETs. We chose to analyze the first three for several reasons. We chose to analyze the first three to balance variance explained from the individual traits with information loss in the calculation of interaction parameters, which involves reparametrizing interaction coefficients from linear models across all traits to new parameters describing the influence of the two loci on each other (equation 3). This recasts all epistasis in terms of two interaction parameters, and therefore introduces a dimensional reduction for three or more ETs. By selecting three ETs, we captured 88.3% of the variance across all eight individual traits, while minimizing information loss in interaction reparametrization. Furthermore, the contributions to the fourth and fifth ETs were primarily from individual traits (Figure 7A), and their inclusion would limit the scope addressed by the final CAPE parameters.

#### Filtering genetic loci for pairwise testing

Because exhaustively testing all marker pairs was computationally infeasible, we filtered markers based on their standardized effects on the ETs. We used linear regression to test the effect of each haplotype at each locus on the ETs and selected the highest-effect haplotypes for pairwise testing. The seven-state linear regression model we used is as follows:

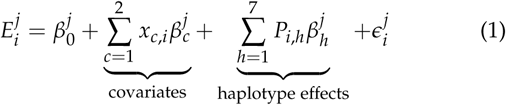

The index *i* is from 1 to number of samples and *j* is from 1 to number of ETs. *P*_*i,h*_ is the probability of each haplotype *h* at the locus, and *x*_*c,i*_ is the presence or absence of each covariate. In this study, we used sex (female:0, male:1) and diet (chow:0, HF:1) as additive covariates.

After calculating the effect of each haplotype on each ET, we filtered the haplotypes to those with the standardized effect size above a minimum threshold for pairwise testing. Retaining all markers in each peak would create large blocks of linked markers, which would inflate the number of selected markers with redundant information. We thus randomly sampled 10% of the markers in each peak along with the marker of maximum effect. We selected a threshold of standardized effect size of 0.11 that yielded 515 individual variants (Supplemental Table S2). Selected haplotypes represented all seven haplotypes across 17 chromosomes and all three ETs. Allele effect sizes were comparable across three ETs, and thus no ET was overly represented by this selection process (Supplemental Figure S1A).

This selection method potentially enriches the sampled marker population for loci with private alleles. This enrichment could be reduced by regressing on SNPs rather than haplotypes, but this procedure would limit interpretation based on genetic ancestry. The method will also miss QTL with interactions that do not yield single-scan main effects, although our permissive acceptance threshold and multiple trait analysis allow mitigate this limitation. Although we enriched the t-statistic distribution for high values, low values were still well represented (Supplemental Figure S1B). Thus, we enriched the marker population for markers most likely to influence on or more traits, but retained the potential to detect many interactions in the absence of significant single-scan main effects.

### Pairwise regression

After filtering the haplotypes to a manageable number, we performed pairwise regression on all pairs of haplotypes as follows:

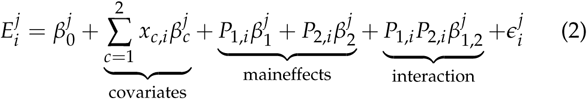

The index *i* runs from 1 to number of samples and *j* runs from 1 to number of ETs. *x*_*c,i*_ is the presence or absence of each covariate. 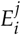 is the ET for sample *i. P*_1, *i*_ and *P*_2, *i*_ are the probabilities of the haplotype at each of two variants for sample *i. P*_1, *i*_ *P*_2, *i*_ is the interaction of two variants, *β*_*1*_ and *β*_2_ are the effects of two variants on ET*j*, and *β*_1,2_ is the interaction coefficient. To avoid testing pairs of closely linked markers, which can lead to false positives, we did not test any marker pair whose Pearson correlation coefficient (*r*) was greater than 0.5.

#### Reparametrization

CAPE coefficients, which indicate the strength and direction of a genetic interaction, are derived from the pairwise regression coefficients through reparametrization. The first step of this process converts the main effect and interaction parameters from the linear regression to two new parameters (*δ*_1_ and *δ*_2_). The *δ* terms can be thought of as the degree to which one variant influences the phenotypic effects of the other. For example, a negative *δ* coefficient indicates that one variant suppresses the effects of the other. If one variant has a negative phenotypic effect, the presence of the other variant suppresses this effect. The *δ* terms are computed in terms of coefficients from pairwise regression as follows:

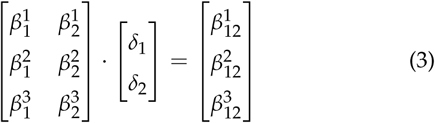

We then translated the *δ* terms into directed variables *m*_12_ and *m*_21_. Unlike the *δ* terms, *m*_12_ and *m*_21_ are directional. They describe how variant 1 influences the effects of variant 2 and vice versa. They are derived from the *δ* terms as follows:

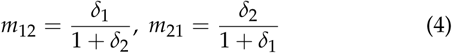

#### Error propagation and significance testing

Each step in our calculations compounds errors in the estimated parameters. We thus propagated the errors through standard least-squares regression and a second-order Taylor expansion on the regression parameters (Carter *et al.* 2012).

We estimated the significance of *m*_12_ and *m*_21_ through permutation testing. Permutations were run by first shuffling the ETs relative to the genotypes and re-running the single-locus regressions and haplotype selection as we did for the true parameter estimation. We then re-ran the pairwise tests on the permuted ETs. We repeated this process until we had generated a null distribution from 500,000 marker pair tests (Tyler *et al.* 2013). We calculated empirical *p* values for each model parameter and corrected these values using false discovery rate (FDR) (Benjamini and Hochberg 1995).

We report the final results in terms of linkage blocks rather than individual markers. We determined block boundaries as described in (Tyler *et al.* 2016). Briefly, for each haplotype, we used the correlation matrix between variants as an adjacency matrix to construct a weighted network, and used the fast greedy community detection algorithm in R/igraph (Csardi and Nepusz 2006) to estimate boundaries between blocks of similar markers.

#### Interpretation of the epistatic network

We analyzed several features of the epistatic network resulting from the CAPE analysis. First, we were interested in the patterns of haplotype interactions. For example, how many interactions were between haplotypes from different founder strains, and how many were between haplotypes from a single founder strain? We were also interested in examining the genetic interactions with the covariates, sex and diet. Genetic interactions with sex could indicate sex-specific risk alleles for metabolic syndrome, while interactions with diet might indicate alleles that either exacerbate or ameliorate the effects of the high-fat, high-sucrose diet.

##### Analysis of variance explained

To estimate the variance explained by covariates, main effects, and interactions, we computed the coefficient of determination for step-wise regression models that included variables for sex, diet, significant main effects, and significant interaction effects added in order. Parameters were only included if significant in the final model.

##### Analysis of network motifs

We assessed the global network architecture. To investigate network structure, we focused on network motifs. We defined network motifs as a pair of interacting haplotypes each with a main effect on a single phenotype (Figure 3). For example, an A/J and CAST haplotype might interact and each influence leptin levels. Motifs can be described as enhancing (positive interaction) or suppressing (negative interaction) and as coherent (both main effects are in the same direction) or incoherent (the main effects are in opposite directions) (Figure 3).

**Figure 3.**
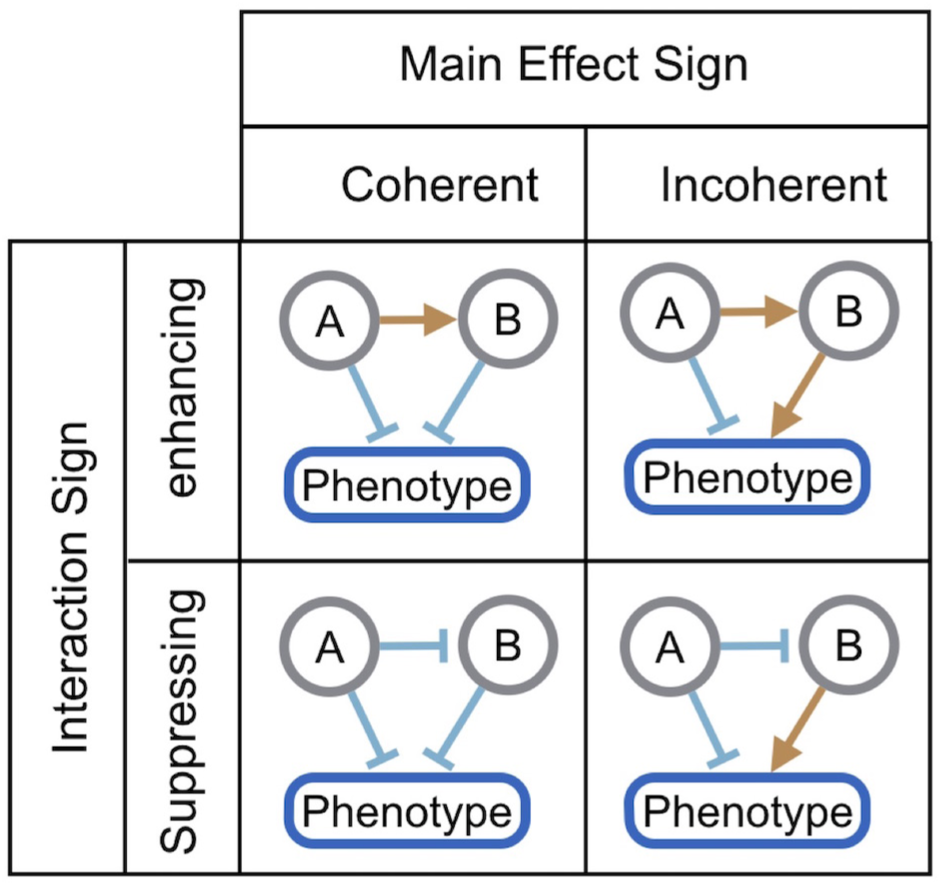
Four types of network motifs. Each motif consists of two markers interacting to influence one phenotype. The markers can either have the same-signed (coherent) or opposite-signed (incoherent) main effects. Their interaction, which can be either enhancing (positive sign) or suppressing (negative sign), may affect additional phenotypes through other main effects.

Previously, we analyzed a large F2 intercross population and found that suppressing-coherent motifs and enhancing-incoherent motifs were enriched in the epistatic network, and that these tended to reduce phenotypic variation in the parental strains (Tyler *et al.* 2016). Thus individuals carrying two reference alleles or two alternate alleles had lower phenotypic variation than individuals carrying one of each at a pair interacting loci. Given the diversity of alternate alleles in this study, we examined these phenotypic effects in the DO epistatic network.

To do this, we identified all the motifs in the DO epistatic network and used linear regression to find the effects of each individual haplotype as well as their interaction effect.

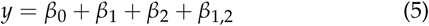

We subtracted the intercept (*β*_0_) from each term (*main*_1_ = *β*_1_ *- β*_0_, *main*_2_ = *β*_2_ *- β*_0_, and *int*_1,2_ = *β*_1,2_ *- β*_0_) and compared the predicted additive effect of the two individual loci (*main*_1_ + *main*_2_) to the actual effect of the full model (*main*_1_ + *main*_2_ + *int*_1,2_).

##### Identification of candidate genes and hypotheses from genetic interactions

Finally, we used information from genetic interactions to generate hypotheses about causal genes in interacting loci. A genetic interaction between loci implies a functional interaction between elements encoded on those loci. We therefore used a function-oriented method to generate hypotheses about which genes in interacting regions might be contributing to the epistatic effects inferred by CAPE. We first identified all protein coding genes in the interacting regions using biomaRt (Durinck *et al.* 2009, 2005). We identified which of these genes had SNPs corresponding to their haplotype effects by querying the Sanger SNP database (Keane *et al.* 2011; Yalcin *et al.* 2011) using the R package SNPTools (Gatti 2015). We further filtered the genes based on functional annotation. We focused on motifs influencing the immune module and thus filtered the genes in each region to those annotated to the Mouse Phenotype (MP) Ontology (Smith *et al.* 2005; Smith and Eppig 2012) term “immune phenotype.” We then looked for the most probable functional interactions between the groups of genes from each chromosomal region using Integrative Multi-species Prediction (IMP) (Wong *et al.* 2015). IMP is a Bayesian network built through integration of gene expression data, protein-protein interaction data, gene ontology annotations and other data. It predicts the likelihood that pairs of genes interact functionally in multiple model organisms and humans. We used IMP to find the highest likelihood connected component that contained at least one gene from each chromosomal region participating in the epistatic interaction. We selected the gene pair with the highest likelihood of interacting functionally as our top candidate gene pair for the interaction.

#### Data Availability

J:DO mice are available for purchase from The Jackson Laboratory (Strain #009376) at https://www.jax.org/strain/009376. Normalized liver gene expression data are available via Gene Expression Omnibus at accession number GSE000000. The physiological phenotypes are described in Supplemental_File_S1_Phenotype_Measurements.rtf, the raw phenotypes are in Supplemental_Table_S1_Phenotypes_Raw.csv and the normalized phenotypes are in Supplemental_Table_S2_Phenotypes_Normalized.csv. The ETs used in the CAPE pipeline are in Supplemental_Table_S3_Phenotypes_Eigentraits.csv, and the filtered genotypes used in the pairwise testing in CAPE are in Supplemental_Table_S4_Filtered_Genotype_File.csv. The complete genotype data for all mice and the R data objects used in all analyses are available at http://do.jax.org. We used the Sanger REL-1505 SNPs and structural variants (Keane *et al.* 2011) and the Ensembl build 82 transcripts (Yates *et al.* 2016). The code used to run the CAPE analysis is in file Supplemental_File_S3_CAPE_DO_code.txt. The code used to run the post-CAPE analyses is in Supplemental_File_S4_cape_additional_scripts.txt. A wrapper for the CAPE analysis is in Supplemental_File_S5_cape_do_analysis.R. A full table of results all interaction and main effect coefficients from CAPE are in Supplemental_Table_S5_Final_Results.csv. The markers included in all linkage blocks are listed in Supplemental_Table_S6_Linkage_Blocks.txt, and the genomic coordinates of all linkage blocks are listed in Supplemental_Table_S7_Linkage_Blocks_Positions.txt. Supplemental_Table_S8_Gene_Lists_for_Expression_Modules.csv contains a list of Ensembl IDs for the genes included in each of the expression modules used in the CAPE analysis. For a full description of supplemental files see Supplemental_Files_Legend.rtf.

## Results

### Transcripts with trans genetic effects cluster into functionally enriched modules

The filtering of the liver transcriptome resulted in 3635 transcripts that were potentially influenced by *trans* genetic loci. The *trans*-eQTL map of these is shown in Figure 4. Effects were broadly distributed, with a potential *trans* hotspot on distal Chr 11 that influences multiple transcripts. We performed WGCNA and obtained 11 modules, three of which were significantly enriched for functional annotations (Huang *et al.* 2009a,b) (Benjamini-adjusted *p* ≤ 0.05). The significant enrichments were: (1) cellular metabolic process (metabolism module) (*p* = 6.3 × 10^-17^), (2) oxidation reduction process (redox module) (*p* = 7.7 × 10^-7^), and (3) immune response (immune module) (*p* = 5.2 × 10^-15^) (Table S1). Modules are referred to hereafter by their functional annotations. We used the three corresponding module eigengenes as phenotypes for CAPE analysis.

**Figure 4.**
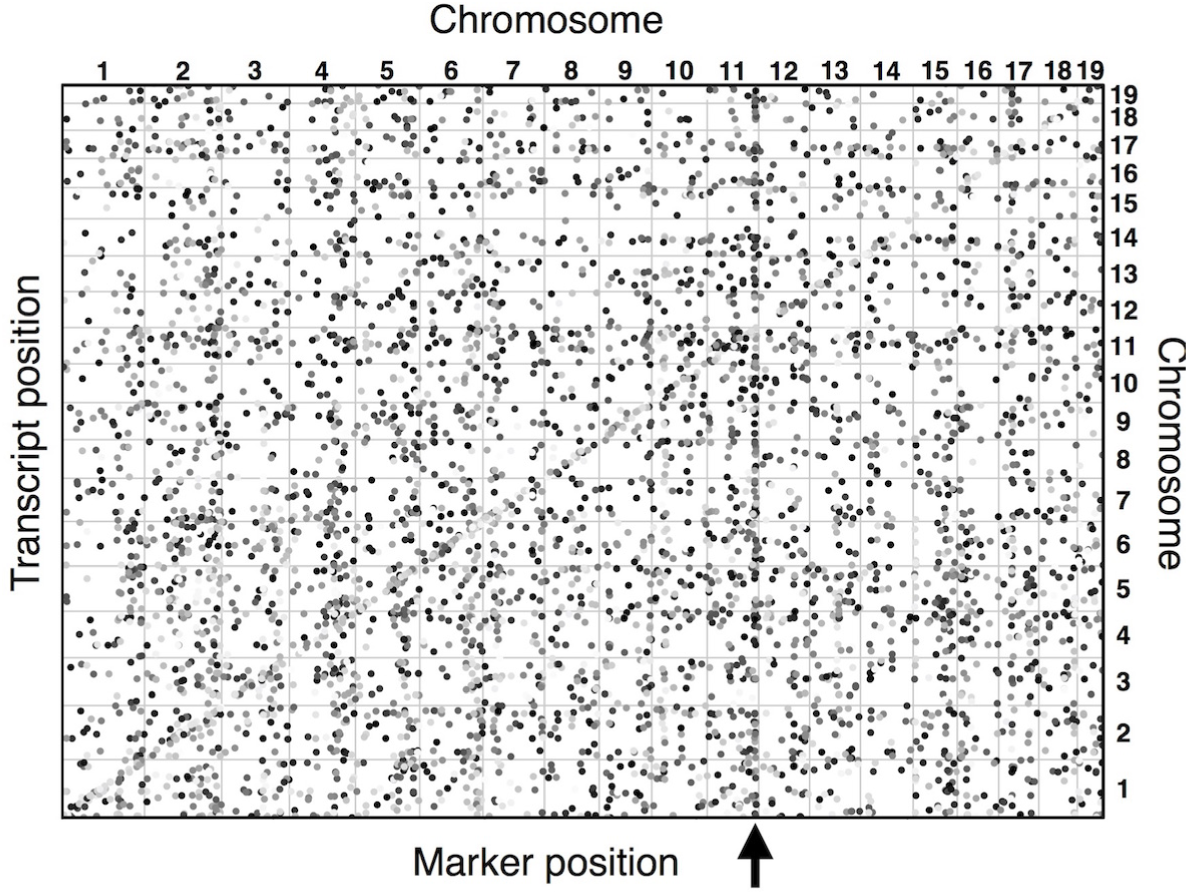
Map of the positions of genes encoding transcripts (y-axis) and their associated eQTL (x-axis). The effect of each eQTL is conditioned on the nearest marker, which eliminates diagonal eQTL likely acting in *cis*. LOD scores range from 7.4 (p = 0.05) to 300, with darker dots represents larger LOD scores. A region on distal Chr 11 (arrow) indicates a potential eQTL hotspot and may encode a gene that influences many transcripts.

### Physiological and expression modules are moderately correlated

The physiological and expression traits in this study were moderately correlated with each other (Figure 5), which implies that there may be common factors (genetic and/or environmental) influencing multiple traits as well as unique information in each phenotype. Correlations (Pearson’s *r*) ranged from -0.54 to 0.75. Fat mass and leptin were the most highly correlated physiological traits (*r* = 0.75, *p* = 2.3 × 10^-86^) and the metabolism module and redox module were the most highly correlated expression traits (*r* = 0.75, *p* = 8.5 × 10^-86^). There were also significant correlations between the physiological traits and the expression traits. The most strongly negatively correlated were lean mass and the metabolism module (*r* = -0.54, *p* = 2.9 × 10^-37^). Triglyceride levels were also negatively correlated with the redox (*r* = -0.36, *p* = 1.3 × 10^-15^) and immune modules (*r* = -0.28, *p* = 1.1 × 10^-9^).

**Figure 5.**
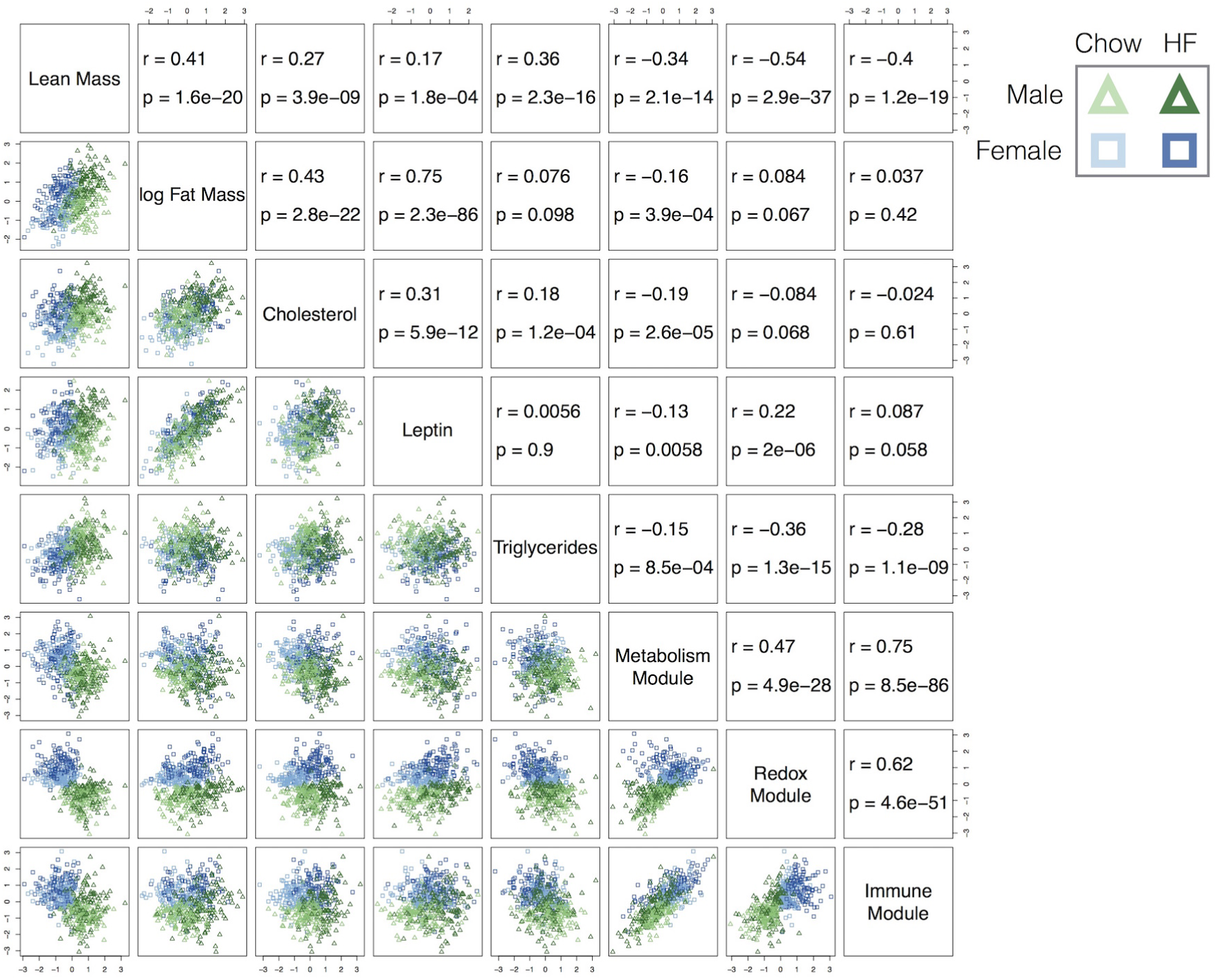
Pearson correlation for all phenotype pairs in this study. Traits tend to be modestly correlated with each other. Physiological traits and expression traits are positively correlated within their groups, but negatively correlated between groups.

### Evidence for pleiotropy influencing physiological and expression traits

The correlations between traits suggest the possibility of pleiotropic loci. Since CAPE uses pleiotropy to infer directionality of interaction effects, we assessed traits for pleiotropy. We first performed a single-locus QTL analysis using DOQTL (Gatti *et al.* 2014). Across all traits, one QTL that influenced cholesterol reached genome-wide significance (permutation-based *p <* 0.01, LOD 8.26) (Figure 6). There were seven additional suggestive QTL (permutation-based p value *p* ≤ 0.63) influencing cholesterol, fat mass, leptin, triglycerides and the metabolism module (See Supplemental Table S10). Two suggestive QTL for fat mass and cholesterol were on distal Chr 11 (Figure 6), potentially indicating a pleiotropic locus. Examination of allele effects at this locus showed a distinct CAST effect on fat mass, cholesterol, leptin, triglycerides, and the redox module (Figure 7). This effect was shared to a lesser extent by the PWK haplotype in fat mass, leptin, and triglyceride levels. As with the LOD scores, these effects did not reach genome-wide significance, but the consistency of the effects is additional evidence of pleiotropy.

**Figure 6.**
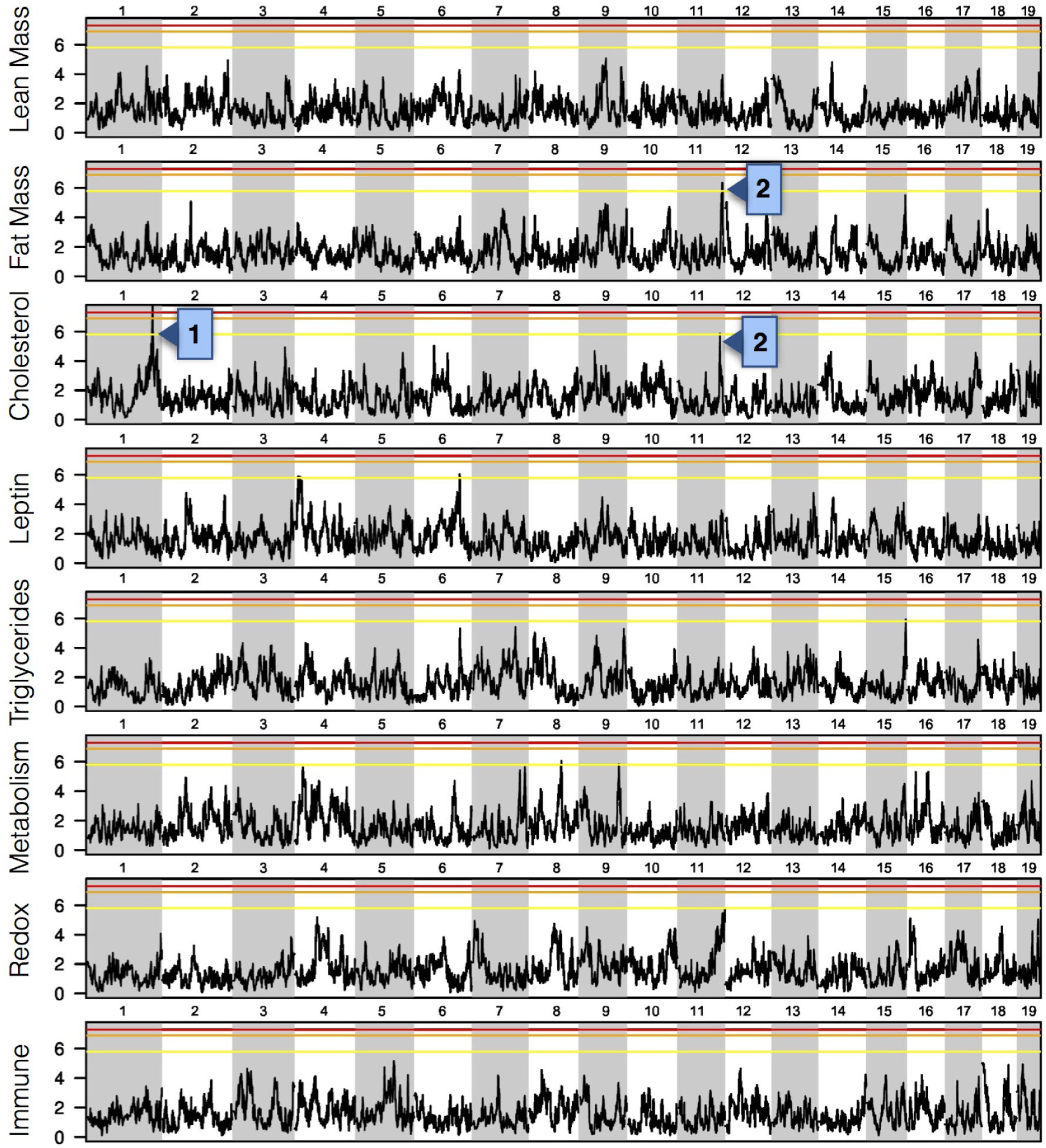
LOD scores for genome scans of all eight phenotypes. A single QTL, at distal Chr 1 for cholesterol, was the only locus at genome wide significance (p < 0.05; Arrow 1). Although no other loci were genome-wide significant, a potentially pleiotropic QTL on distal Chr 11 is suggestive for both fat mass and cholesterol (Arrow 2). Horizontal lines denote permutation-based thresholds of p < 0.05 (red, LOD 7.3), p < 0.01 (orange, LOD 6.9), and p < 0.63 (yellow, LOD 5.8).

In addition to pleiotropic effects, CAPE requires unique genetic effects across multiple traits to provide non-redundant information for inference of epistatic interaction directionality. There was evidence for unique effects across the genome for all traits. For example, the NZO haplotype on distal Chr 11 had a positive effect on cholesterol and the A/J haplotype had a positive effect on leptin (Figure 7).

**Figure 7.**
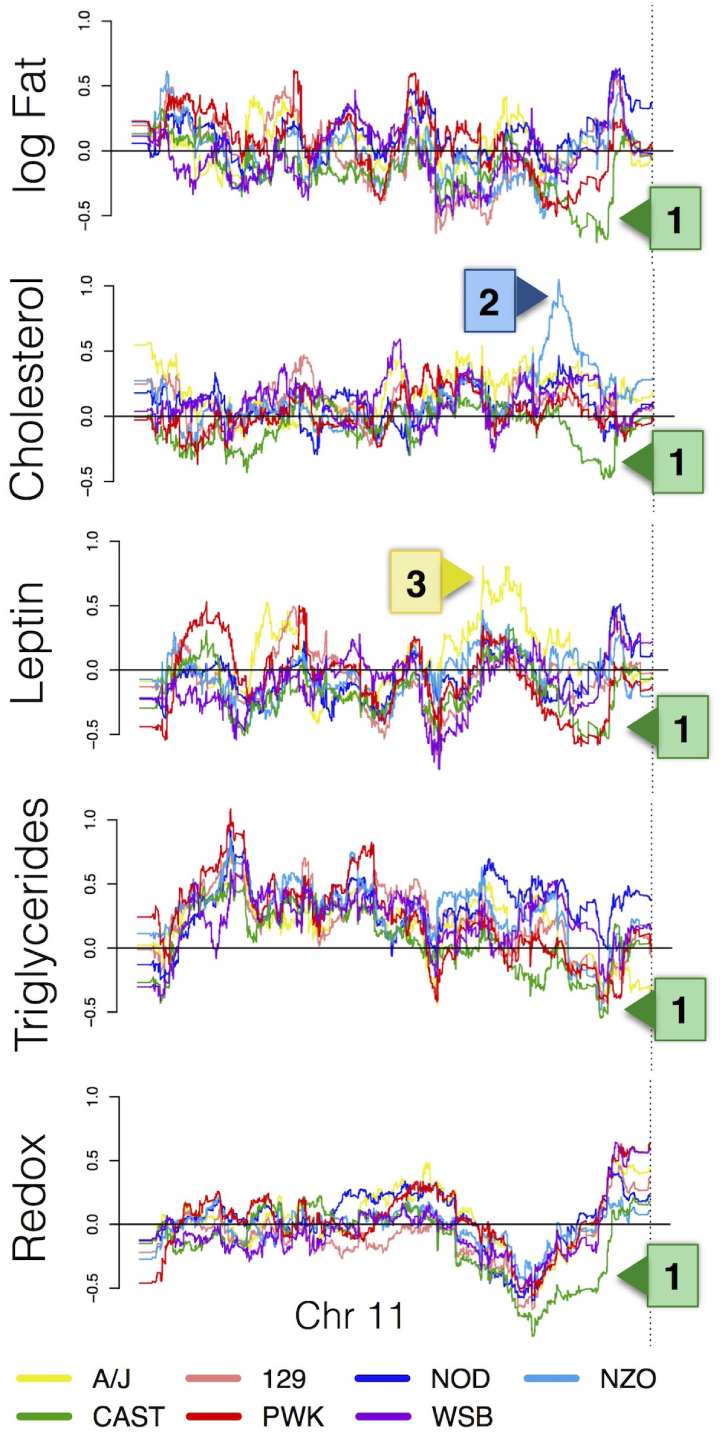
Estimated allele effects of each strain haplotype on Chr 11 for five traits: log fat mass, cholesterol, leptin, triglycerides, and the redox expression module. The CAST haplotype on distal Chr 11 has pleiotropic effects on all traits (Arrow 1). The NZO and A/J haplotypes have individual effects on cholesterol (Arrow 2) and leptin (Arrow 3), respectively. All effects are relative to the B6 reference.

### Singular value decomposition concentrates pleiotropic effects

We used singular value decomposition (SVD) to decompose the trait matrix into eight orthogonal eigentraits (ETs) (Materials and Methods; Figure 8A). This procedure recombines covarying elements of the measured traits, and potentially concentrates functionally related effects making them easier to detect. In our subsequent analysis, we used the first three ETs, which captured 88.3% of the overall variance. The fourth and fifth ET primarily represented cholesterol and triglyceride phenotypes, respectively, and their inclusion in CAPE modeling biases models to fit these variances at the expense of other traits. DOQTL identified a single locus on distal Chr 11 that influenced ET2 (Figure 8B) (permutation-based *p* ≤ 0.1). From the analyses of the individual traits, we had identified this region as potentially pleiotropic, influencing both fat mass and cholesterol, but the locus did not achieve genome-wide significance in either trait. The greater significance for ET2 suggests that ET2 aggregates signals from fat mass and cholesterol that are influenced by a pleiotropic locus on Chr 11. This was further supported by the allele effects, in which CAST showed the greatest influence (Figure 8C). Interestingly, there were no other significant or suggestive QTL for the ETs.

**Figure 8.**
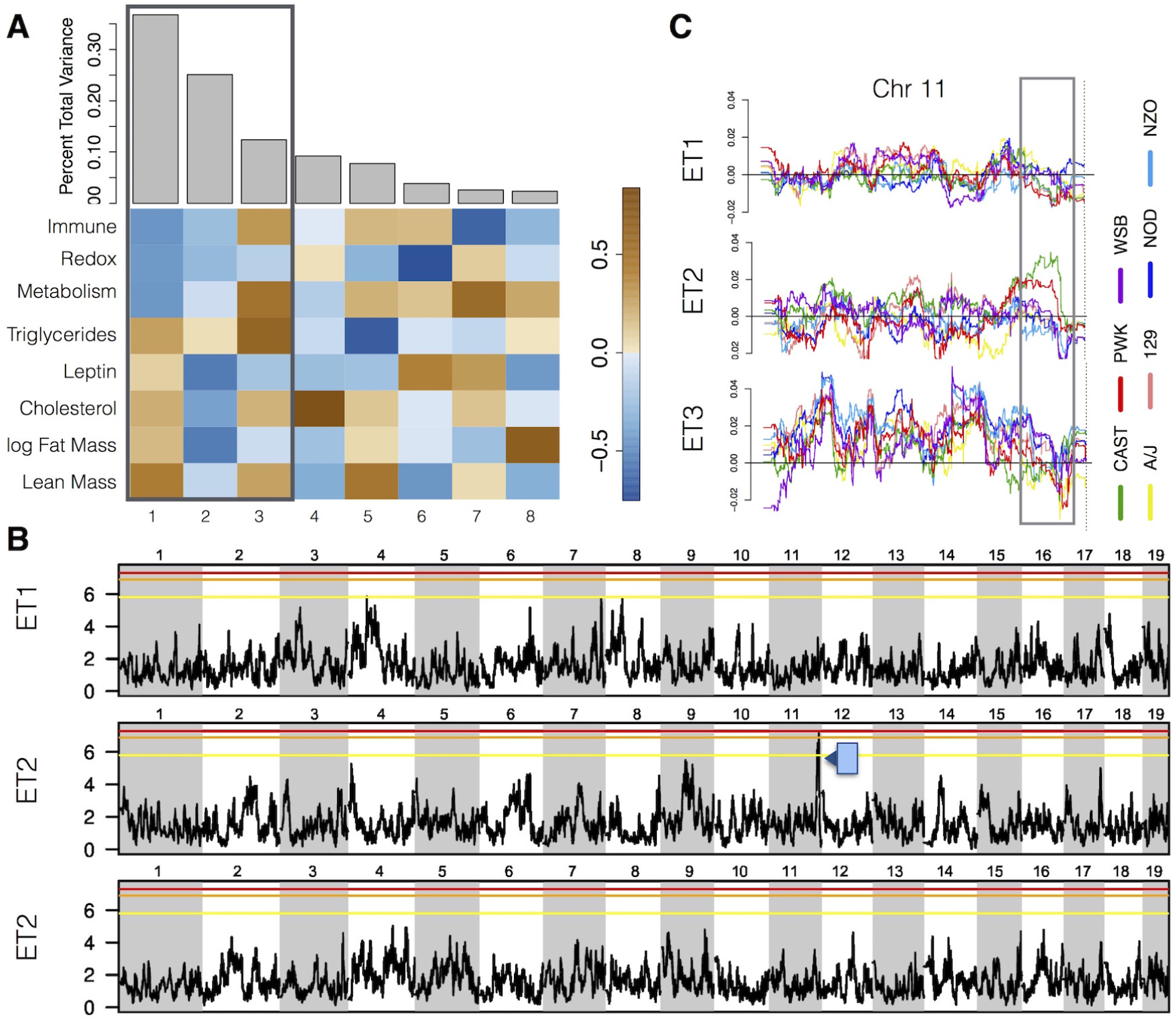
Eigentraits (ETs) of combined phentoypes determined by singular value decomposition. A) Eight orthogonal ETs and their varaince content. Proportion of the total variance captured by each ET (gray bars) and relative contributions of each trait to each ET (heatmap) are shown. The box highlights the three ETs selected for CAPE analysis. B) LOD scores for genome scans of the first three ETs. One QTL on distal Chr 11 was suggestive (p < 0.2, arrow) and may reflect a pleiotropic locus influencing both fat mass and cholesterol. The red and black horizontal lines are at permutation-based thresholds of p < 0.05 (LOD 7.49) and p < 0.2 (LOD 6.54), respectively. C) Individual haplotype effects for Chr 11 on the first three ETs. The box highlights the QTL location for ET2.

### An epistatic network involving all haplotypes influences physiological and expression traits

After calculating haplotype effects on the three ETs, we sampled 515 individual haplotypes with the largest effect sizes across the genome for pairwise testing (Materials and Methods). We used CAPE to test these markers for directed epistatic interactions affecting the first three ETs (Materials and Methods). The resulting network consisted of 89 significant interactions among 49 loci and two covariates (Figure 9). Significance was based on 500,000 permutations and set at an FDR-adjusted q value *≤* 0.05. To determine the robustness of these interactions, we bootstrapped the standardized effects by sampling the 474 mice in the experiment with replacement and re-running the CAPE pipeline. Both main effect (Supplemental Figure S2) and interaction (Supplemental Figure S3) standardized coefficients were robust to sampling.

**Figure 9.**
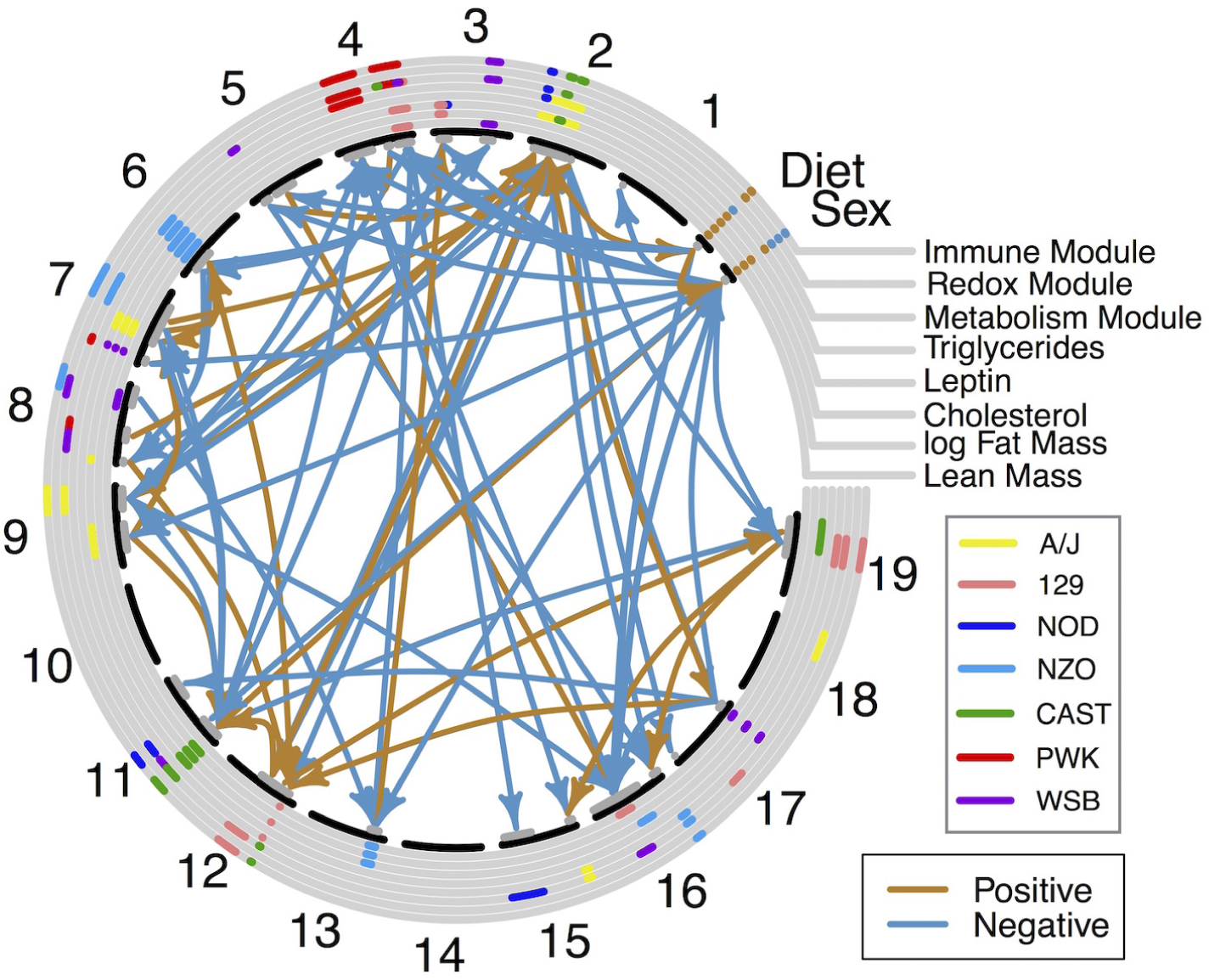
The pleiotropic QTL interaction network derived from CAPE. Main effects, colored by haplotype, appear in the concentric circles. Sex and diet main effects are shown in brown or blue to denote positive and negative effects, respectively. Arrows between chromosomal regions denote genetic interactions that indicate an enhancing effect (brown) or a suppressing effect (blue).

For each trait, we calculated the variance explained by the significant main effects, the significant interactions, and by each of the covariates (Figure 10). The total variance explained ranged from 23% for triglycerides to 66% for the redox expression module. For all traits, the covariates, sex and diet, accounted for the majority of the variance explained. Main effect loci explained an additional 13% and 19% variance in lean mass and the metabolism module, respectively. Relatively little additional variance was explained by interactions. The largest proportions were explained in lean mass and leptin which gained a further 1.8% and 1.6% variance explained, respectively.

**Figure 10.**
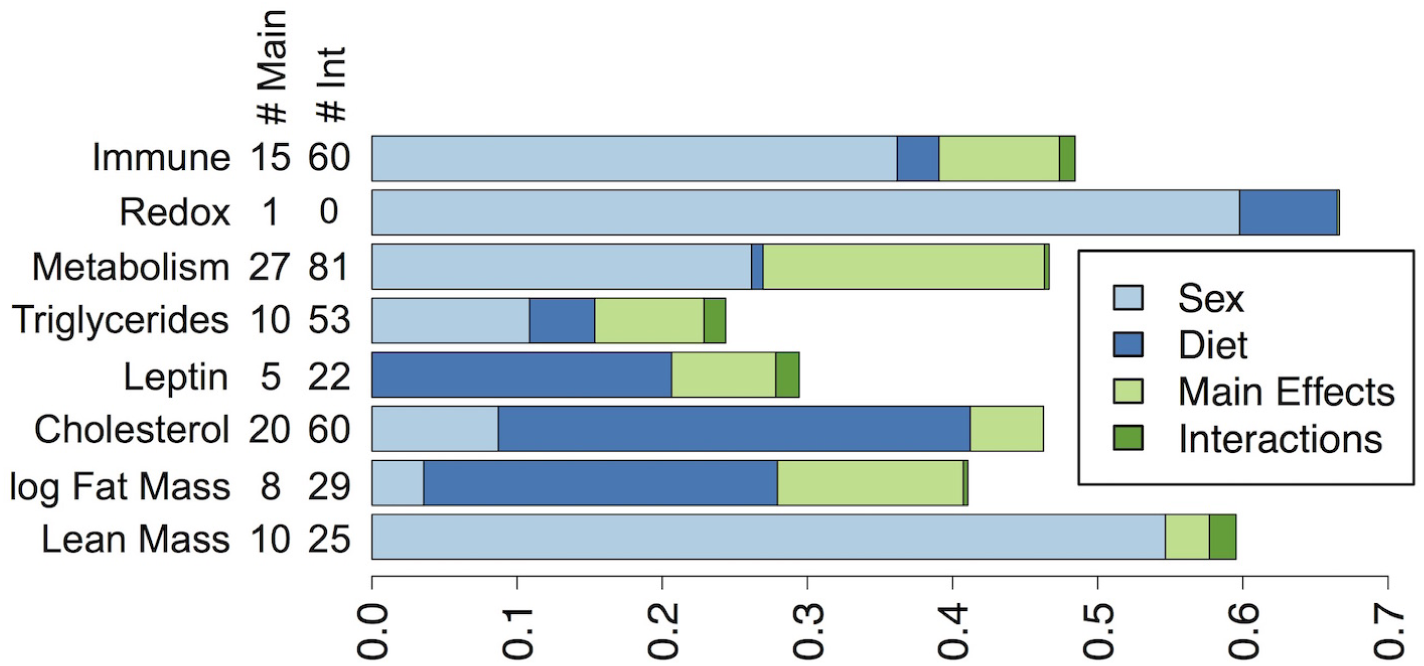
Variance explained by covariates, main effects, and interactions. Bars show the proportion of variance of each trait explained by the covariates, sex (light blue) and diet (dark blue), the main QTL effects (light green), and by the CAPE-derived genetic interactions (dark green). Numbers in the left margin report the number of significant main effects (# Main) and the number of significant interactions (# Int) influencing each trait.

To determine strain contributions to this network, we assessed how frequently individual haplotypes were involved in genetic interactions. We found that each haplotype participated in at least one interaction (Figure 11A). WSB alleles were involved in the greatest number of interactions (32), while NZO alleles participated in the fewest (8). The total number of interactions for haplotype was marginally correlated with its representation in the 515 markers selected for analysis (Figure 11A) (*p* = 0.1). Most haplotypes were balanced in terms of source and target in the directed interactions. However, the 129 haplotype was a target of interactions about four times more frequently than it was a source, while the NZO haplotype was a source about twice as many times as it was a target (Figure 11A). The covariates, sex and diet, were both much more frequently sources of interactions than they were targets (Figure 11A), suggesting that these factors commonly modify genetic effects and that their broad effects are less commonly adjusted by genetic factors.

**Figure 11.**
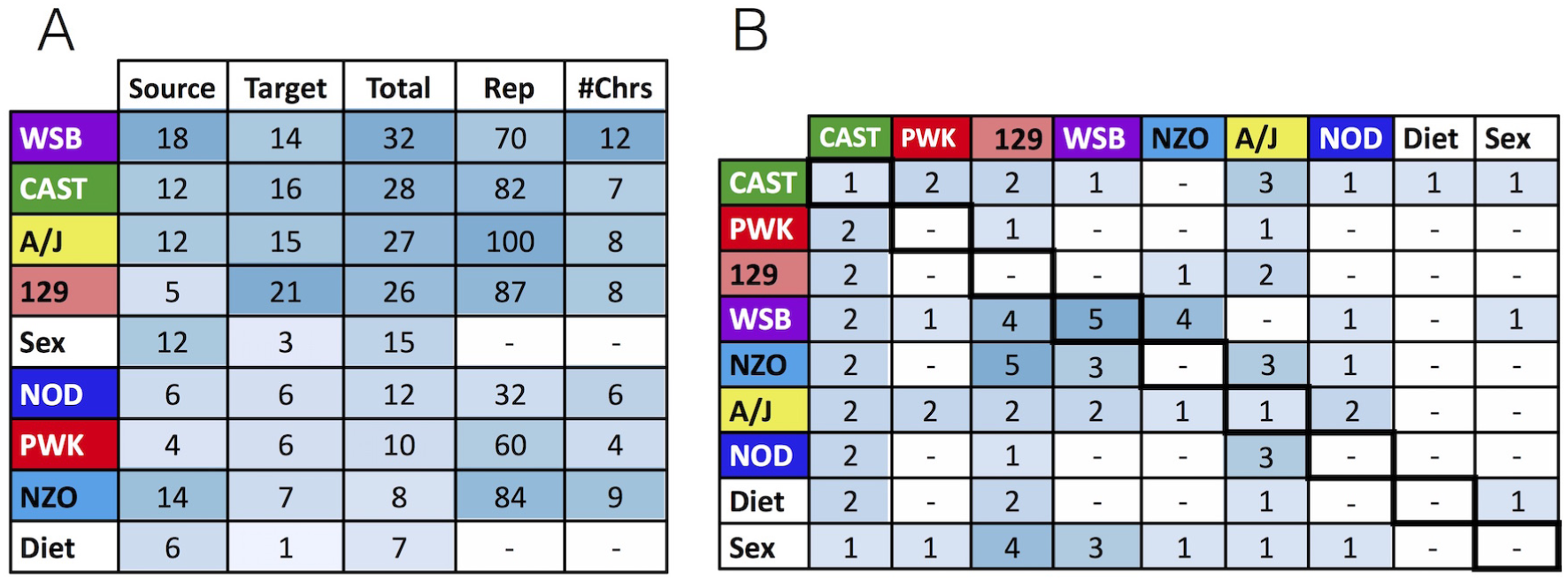
Frequency of haplotype participation in genetic interactions. A) The number of times haplotypes of each ancestry was the source or target of an interaction, sorted by total number of interactions. The final two columns indicate how many candidate markers were tested for pairwise interactions, and the total number of chromosomes containing the markers. Shading highlights higher counts. B) A detailed count of the interactions by ancestry and covariate. Darker squares represent higher counts, and counts of 0 are represented by dashes for clarity.

We next determined how frequently haplotypes interacted in order to compare cross-strain and within-strain interaction frequencies (Figure 11B). For the seven founder haplotypes, there were seven possible intra-haplotype interactions and 21 possible inter-haplotype interactions. We found seven intra-strain interactions and 61 inter-strain interactions. This excess of crossstrain interactions (nine time more common than intra-strain interactions or three times expectation) suggests functional mismatches between haplotypes are more frequently observed than the recovery of subnetworks private to a single founder. Five of the seven intra-strain interactions were between WSB alleles, which is potentially due to the greater number of WSB markers meeting the testing criteria. Inter-strain interactions were concentrated among 129, WSB, NZO and A/J alleles, which are all members of the *Mus musculus domesticus* subspecies. CAST, *M. musculus castaneus*, interacted with each of the other strains relatively evenly, while PWK, *M. musculus musculus*, was the most isolated strain, and did not interact at all with the NZO or NOD haplotypes.

### Epistatic network hub overlaps trans-eQTL hotspot

A minority of QTL were involved in multiple genetic interactions. The genetic locus participating in the largest number of interactions (nine) was a CAST haplotype on Chr 11. This haplotype (98.2 Mbp to 117.5 Mpb) coincided with the pleiotropic CAST haplotype influencing fat mass, cholesterol, leptin, triglycerides, and the redox module (Figure 7), as well as an apparent trans-eQTL hotspot (Figure 4). Taken together, these results suggest that the multiple traits influenced here are related to each other through redox related transcriptional programs, and that this locus may contain a master regulator that influences blood lipid profiles and body composition in part through redox related transcription.

### Sex interacted with diet and QTL from all founder haplotypes

We observed multiple interactions between sex and QTL, suggesting that sex modifies genetic effects or, conversely, some genetic effects are sex-specific. As previously observed, sex had significant effects on all physiological traits except leptin levels (Eppig *et al.* 2015). This effect was positive for all phenotypes, such that males had higher log fat mass (males 1.9 g, females: 1.7 g, *p* = 5.7 × 10^-2^), lean mass (males: 25.1 g, females: 18.3 g, *p <* 2 × 10^-16^), cholesterol (males 110.4 mg/dl, females: 93.8 mg/dl, *p* = 4.3 × 10^-10^), and triglycerides (males: 156.0 mg/dl, females: 115.0 mg/dl, *p* = 7.6 × 10^-14^). All expression modules were significantly lower in males (all *p <* 2 × 10^-16^).

The majority of genetic interactions with sex (12 of 15) involved a suppression of allele effects by sex, indicating that the alleles had larger effects in females than in males. At least one QTL from each founder strain was affected. In three cases, the effects of sex were modified by a QTL or high-fat diet. Since these are cases in which a QTL broadly affected many traits via its influence on all sex effects, we consider them in more detail. First, a CAST allele at a Chr 11 QTL enhanced the effects of sex, illustrated with the metabolism module (Figure 12A). Both the CAST allele and male sex reduced this module phenotype, and the joint effect was greater than the additive expectation. Thus these factors act synergistically to affect the module. The second interaction is a WSB allele on Chr 17 that suppresses sex effects, and can be illustrated with cholesterol as a phenotype (Figure 12B). Both the WSB allele and the male sex increase cholesterol levels, but the joint phenotype is lower than expected from an additive model due to the QTL reducing the sex effect. Finally, phenotypic effects of male sex were enhanced by diet, suggesting that males were more susceptible to the effects of a high-fat (HF) diet (Figure 12C). The HF diet slightly reduced triglyceride levels, while males had greater triglycerides than females. However, males on the HF diet had greater triglyceride levels than expected from the additive model. Diet enhanced the positive effect of the male sex, overriding the negative marginal effect of diet.

**Figure 12.**
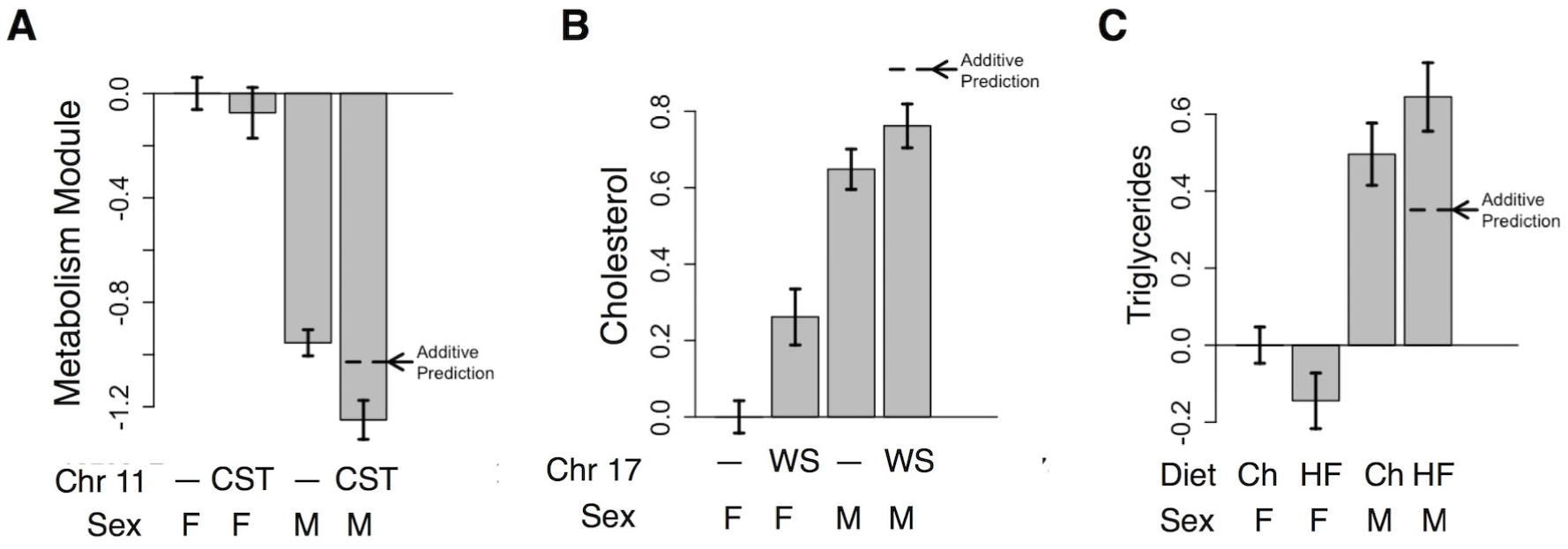
Examples QTL-sex and sex-diet interactions. A) The CAST (CST) haplotype at a Chr 11 QTL had a negative effect on the metabolism module relative to all non-CAST haplotypes (—), and interactively enhanced the effect of the male sex. Thus in males with the CAST haplotype, the metabolism module was lower than expected from the additive model. B) The WSB (WS) haplotype at a Chr 17 QTL had a positive effect on cholesterol relative to all non-WSB alleles (—). Males also have higher cholesterol than females. The WSB allele suppressed this effect in males, however, and males with the WSB allele had lower cholesterol than expected from the additive model. C) The high fat diet (HF) had a negative effect on triglyceride levels relative to the chow diet (Ch), and males had higher triglycerides than females. However, males on the high-fat diet had higher triglyceride levels than expected from the additive model. Diet enhanced the positive effect of the male sex (M) which, in turn, overcame the negative marginal effect of diet. In panels A and B bars show mean phenotype values for animals partitioned by sex and genotype. In panel C bars show phenotype means for animals partitioned by sex and diet. Error bars denote standard errors.

### Diet interacted with QTL from a subset of founder haplotypes

In addition to its interaction with sex, diet also interacted with multiple genetic loci. On its own, diet significantly increased log fat mass (chow: 1.6 g, HF: 2.1 g, *p <* 2 × 10^-16^), cholesterol (chow: 85.8 mg/dl, HF: 119.1 mg/dl, *p <* 2 × 10^-16^), and leptin (chow: 7.7 mg/dl, HF: 19.7 mg/dl, *p <* 2 × 10^-16^) and significantly decreased triglyceride levels (chow: 146.7 mg/dl, HF: 124. 3 mg/dl, *p* = 1 ×10^-4^). It also significantly decreased all expression modules (all *p <* 0.001). Similar to sex, the majority of genetic interactions with diet (five of seven) were those in which high-fat diet suppressed genetic effects. That is, the genetic loci had greater phenotypic effect in chow-fed mice than mice on the high-fat diet. These QTL may therefore mimic a high-fat diet in their effects on their targeted phenotypes. There was one locus, the CAST allele on Chr 2, that enhanced the effects of diet, indicating that animals carrying this haplotype were more susceptible to all effects of the high-fat diet. The effects of diet were also enhanced by sex, as mentioned above, indicating that males in this population were more susceptible to the effects of the high-fat diet than females.

### Network motifs reveal both redundant and synergistic genetic interactions

To better understand the overall influence of genetic interactions on traits, we performed network motif analysis (Tyler *et al.* 2016). Network motifs were defined as one interaction between two loci that each had a main effect on one phenotype (Figure 3). The interaction was either suppressing or enhancing, and the two main effects affected the phenotype in either the same direction (coherent) or opposing directions (incoherent). Here we investigated the effects of network motifs on traits in the DO and compared the results to the F2 intercross in Tyler *et al.* (2016). Only enhancing-incoherent and suppressing-coherent motifs were present in the DO epistatic network (Figure 13). These were the most common class of interactions in the F2 intercross, in which instances of both motifs tended to reduce trait variability in founder strains. That is, animals with the same parental haplotype at two interacting loci had lower trait variation than animals with one of each parental haplotype. Thus haplotypes from different strains had incompatible effects, destabilizing traits by driving them to extreme values (Tyler *et al.* 2016). Here we found that in the DO population 70% of suppressing-coherent interactions tended to stabilize traits as they did in the F2 intercross. On the other hand, 72% of enhancing-incoherent interactions had synergistic effects and tended to destabilize traits. A substantial fraction (28%) of enhancing-incoherent interactions had extreme synergistic effects, driving traits past additive predictions from any QTL pair (Figure 14). Less than 1% of the suppressing-coherent motifs had this effect. In a two-parent cross there is only one alternative strain, so epistasis denotes interaction between two alternative alleles derived from the same parent. In contrast, a multiparental population like the DO enables epistasis between alleles from multiple founder lines. The majority of interactions involved alleles from different founders for both the enhancing-incoherent (72%) and suppressing-coherent (96%) motifs.

**Figure 13.**
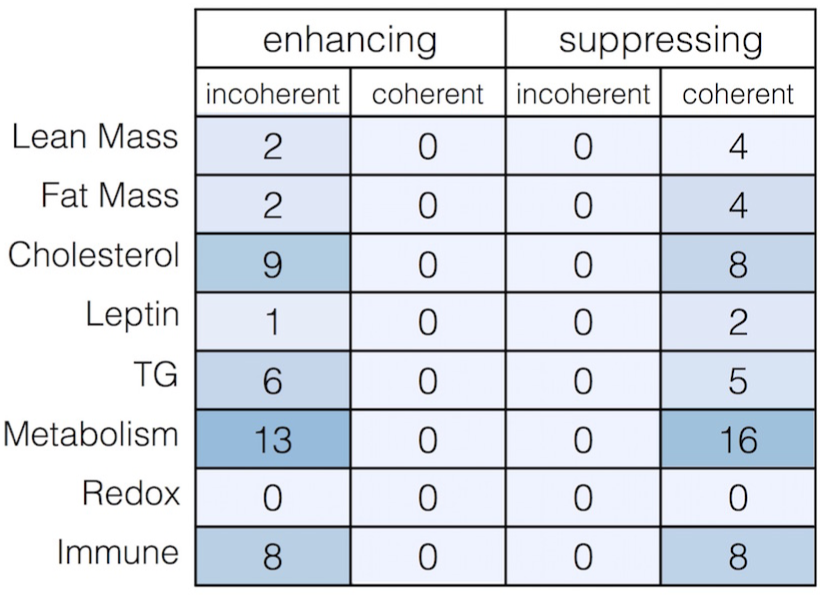
Counts of each different motif type in the QTL-QTL network for each phenotype. Darker shading indicates higher counts.

**Figure 14.**
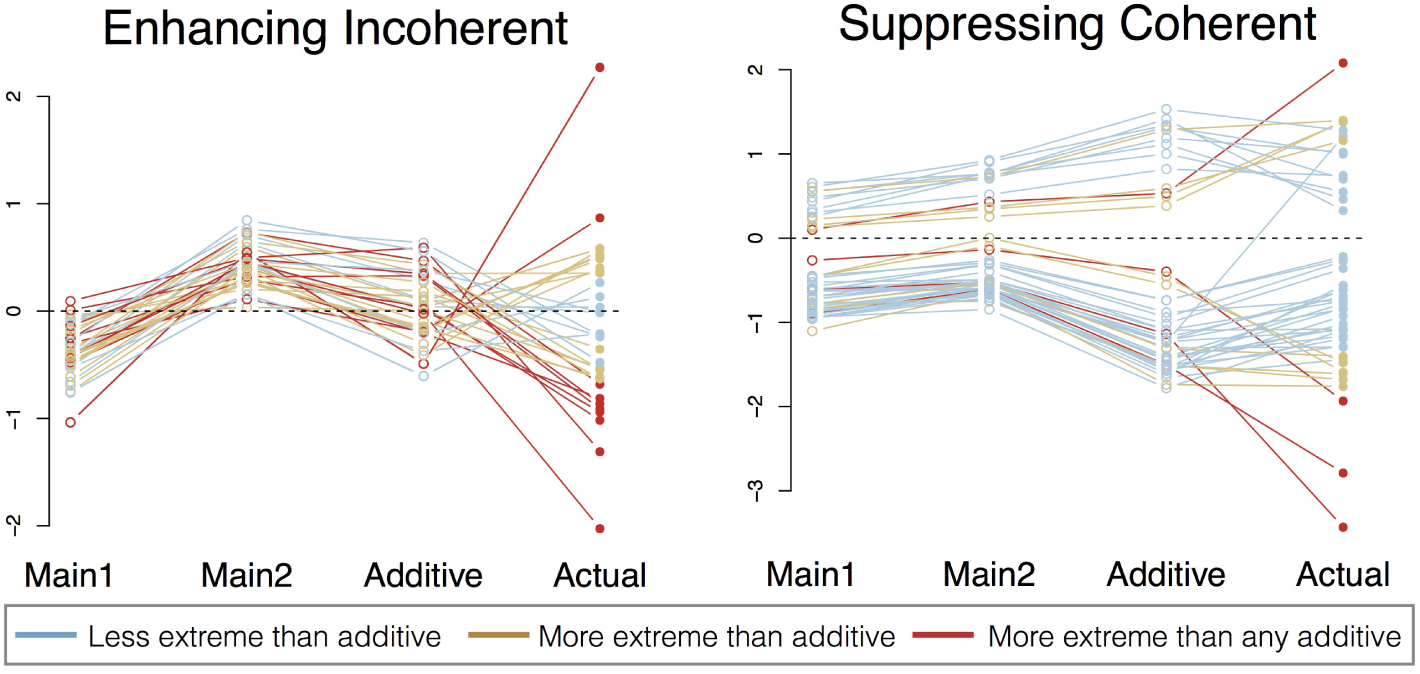
Phenotypic effects of enhancing-incoherent (left) and suppressing-coherent (right) network motifs. Main1 and Main2 denote the average deviation from population mean in normalized phenotype for animals carrying the alternate haplotype at the two QTL. Marker 1 and marker 2 are sorted such that marker 1 always has the lesser effect. Additive is the predicted additive effect determined by the sum of Main1 and Main2. Actual is the observed deviation from the population mean of animals carrying the alternate haplotype at both markers. Lines are drawn to connect dots from individual interactions. Blue and brown lines indicate motifs that bring phenotypes closer and further to the population mean than predicted by the additive model, respectively. Red lines indicate a subset of motifs that exhibit phenotypes more extreme than would be predicted by any additive model.

## Discussion

Traits related to metabolic disease, such as cholesterol levels and body fat mass, have complex genetic architecture. Here we used Combined Analysis of Pleiotropy and Epistasis (CAPE) to identify an epistatic network influencing of body composition, serum biomarkers, and hepatic gene expression in DO mice. The network linked genetic loci to each other, as well as to sex and diet, providing an overview of the complexity of these related traits. The network also serves a scaffold that can be used to generate specific hypotheses about genes influencing individual traits.

Although our study is likely underpowered, our CAPE analysis found that epistasis is abundant in DO mice. Pervasive epistasis has been observed in many different organisms and in different experimental paradigms (Mackay 2014) including flies (Horn *et al.* 2011), nematodes (Lehner *et al.* 2006; Byrne *et al.* 2007), mice (Pavlicev *et al.* 2010; Shao *et al.* 2008), yeast (Tong *et al.* 2001; Segrè *et al.* 2005; Snitkin and Segrè 2011), maize (Ma *et al.* 2007), and arabidopsis (Rowe *et al.* 2008). CAPE has been previously used to identify epistatic networks in mice (Tyler *et al.* 2016), *Drosophila* (Carter 2013), and yeast (Carter *et al.* 2012). From these studies, we found that the structure of the networks across organisms is similar, with pervasive epistasis and balanced numbers of suppressing and enhancing interactions. Although varying definitions of epistasis make direction comparision between studies difficult, both positive epistasis and negative epistasis have been similarly detected in systems ranging from yeast (Tong *et al.* 2001; Segrè *et al.* 2005) to mammalian models (Pavlicev *et al.* 2010).

The interaction effects in the epistatic network tended to be weak (standardized effect mean: 2.18, sd: 0.4) relative to main effects (standardized effect mean: 4.55, sd: 0.47), and account for only marginal trait variance (Figure 10). This is consistent with epistasis previously found in intercross and outbred populations (Bloom *et al.* 2015; Shao *et al.* 2008; Mackay and Moore 2014; Mackay 2014). However, this does not imply that the interactions are negligible as they may be critical in predicting phenotypes in individuals (Forsberg *et al.* 2016; Mackay and Moore 2014; Shao *et al.* 2008; Nadeau and Dudley 2011; Nadeau 2003). For example, genetic interactions explained only 1.0% of the variance of the immune module when averaged across the entire population. However, using additive models to predict immune module levels in animals with epistatic alleles leads to large errors. In animals carrying the CAST allele on Chr 2 and the A/J allele on Chr 9, immune module levels are dramatically higher than predicted by the additivity (Figure 15A). As shown in the motif analysis (Results), many of the genetic interactions in the network result in extreme trait values. Extending these observations to humans, identifying individual genetic interactions may be critical to predicting disease risk in individuals (Mackay and Moore 2014; Nadeau and Dudley 2011; Nadeau 2003) even if they explain relatively little trait variance across the population (Hill *et al.* 2008).

**Figure 15.**
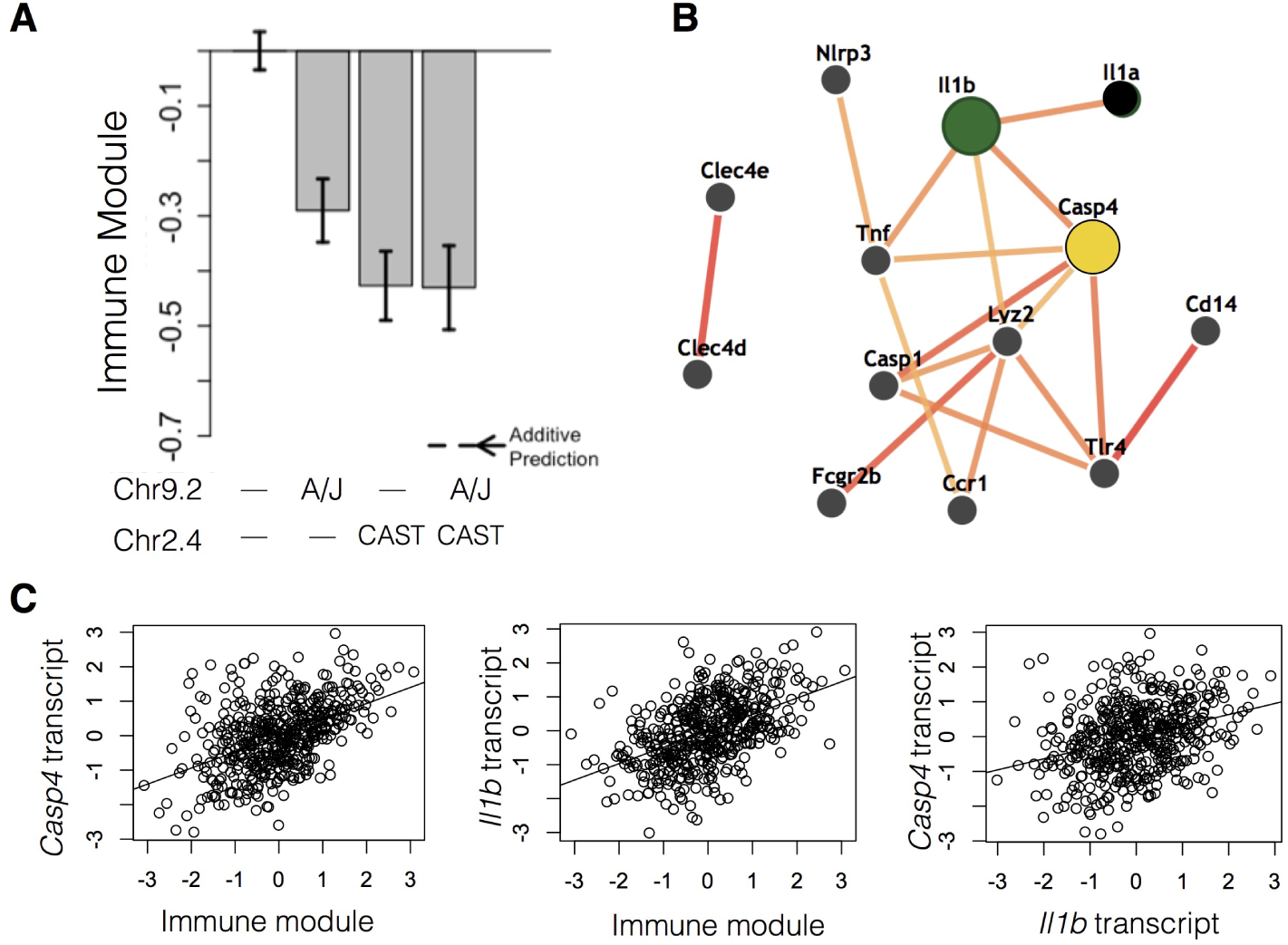
Gene prioritization for interacting QTL Chr9.2 and Chr2.4. A) Both the A/J haplotype at Chr9.2 and the CAST haplotype at Chr2.4 have negative effects on the immune module. Together, they have an effect similar to that of the CAST haplotype at Chr Error bars show standard error. B) Functional connections between Il1b and Casp4 from the IMP network. The two proteins are predicted to interact functionally with high confidence. C) The transcripts of *Casp4*, in Chr9.2, and *Il1b*, in Chr2.4, are both correlated with the immune module. The transcripts are also correlated with each other.

Our network motif analysis highlighted the importance of identifying individual epistatic interactions in outbred populations. We found that suppressing-coherent motifs tended to reflect redundancy, while the enhancing-incoherent motifs often drove traits to extreme values. This is in contrast to our previous findings in a mouse intercross (Tyler *et al.* 2016), in which both types of motifs tended to stabilize traits near the population mean. This is potentially due to within-strain accumulation of alleles to maintain trait homeostasis; as more same-strain alleles are combined, the trait is stabilized. In contrast, the phenotype destabilization we observed in the DO may be related to novel haplotype combinations from the eight founders. Novel recombinations of alleles from multiple strains may combine incompatible molecular strategies for maintaining homeostasis and drive traits to the extreme values we observed. Alternatively, it has been observed that genetic architecture can be trait-specific (Shao *et al.* 2008; Snitkin and Segrè 2011), and it is possible that the difference in the traits analyzed in the DO and intercross studies accounts for the differences in genetic architecture.

While identification of genes in QTL is challenging due to multiple positional candidates, the generation of molecular hypotheses can augmented by combining the functional information in epistasis with gene annotations. We chose two candidate interactions for examples of hypothesis generation. First, we considered a suppressive interaction between the A/J haplotype on Chr 9 locus 2 (Chr9.2: 5-36 Mb) and the CAST haplotype on Chr 2 locus 2 (Chr2.2: 123-133 Mb). Both haplotypes had a negative main effect on the immune module, and their combined effect exhibited genetic redundancy (Figure 15). Data integration (Materials and Methods) identified *Casp4* in Chr9.2 and *Il1b* in Chr2.2 as the most likely candidate genes. Supporting this hypothesis, the abundance of both transcripts are correlated with the immune module (*Casp4*: *r*^2^ = 0.48, *p* = 2.6 × 10^-28^, *Il1b*: *r*^2^ = 0.49, *p* = 1 × 10^-30^), and with each other (*r*^2^ = 0.32, *p* = 7.4 × 10^-13^) (Figure 15C). *Casp4*, also known as *Casp-11*, is a member of the cysteine-aspartic acid protease family and is essential for IL1B secretion, and mice with homozygous mutations of *Casp4* have decreased levels of circulating IL1B (Wang *et al.* 1998). These findings are consistent with the redundant genetic interaction we observed between Chr9.2 and Chr2.2. Redundant interactions are hypothesized to occur between variants encoding genes within a single pathway (Avery and Wasserman 1992; Lehner 2011). Each variant had a similar effect on the pathway but, because pathway function can only be disrupted once, the combination of the two variants did not have a further effect despite inheritance from different founder strains.

Next, we examined an enhancing interaction between the same A/J haplotype in Chr9.2 and a distinct QTL on Chr 2. This second locus, Chr 2 locus 4 (Chr2.4: 165-171 Mb) represented an effect of the NOD haplotype and did not overlap the CAST-driven QTL Chr2.2 above. The A/J Chr9.2 and the NOD Chr2.4 loci influenced the immune expression module in opposite directions, and together, they drove the trait to be slightly more negative than predicted by the additive model (Figure 16A). Our gene prioritization identified *Casp4* again for the Chr9.2 A/J locus, and *Src* as a likely interacting partner in the Chr2.4 NOD locus (Figure 16B). Transcripts for both genes are significantly correlated with the immune module (*Casp4*: *r* = 0.47, *p* = 6.3 × 10^-28^; *Src*: *r* = 0.47, *p* = 3.7 × 10^-27^) and with each other (*r* = 0.21, *p* = 3.2 × 10^-6^) (Figure 16C). In the IMP network *Casp4* and *Src* occupy two lobes of a connected graph, suggesting that they are less directly functionally related than *Casp4* and *Il1b*. The *Casp4* region of the network is enriched for genes involved in inflammasome pathways (*p* = 2.9 × 10^-6^) (Motenko *et al.* 2015), while the *Src* subnetwork is enriched for EGFR signaling (*p* = 2.7 × 10^-4^) (Motenko *et al.* 2015). The IL-1 and EGF families of proteins are upregulated in human keratinocytes during wound healing and in psoriasis, and they have been shown to interact synergistically in upregulating transcripts involved in antimicrobial defenses (Johnston *et al.* 2011). These observations suggest that the A/J allele of *Casp4* and the NOD allele of *Src* may interact to influence immune-related expression in mice.

**Figure 16.**
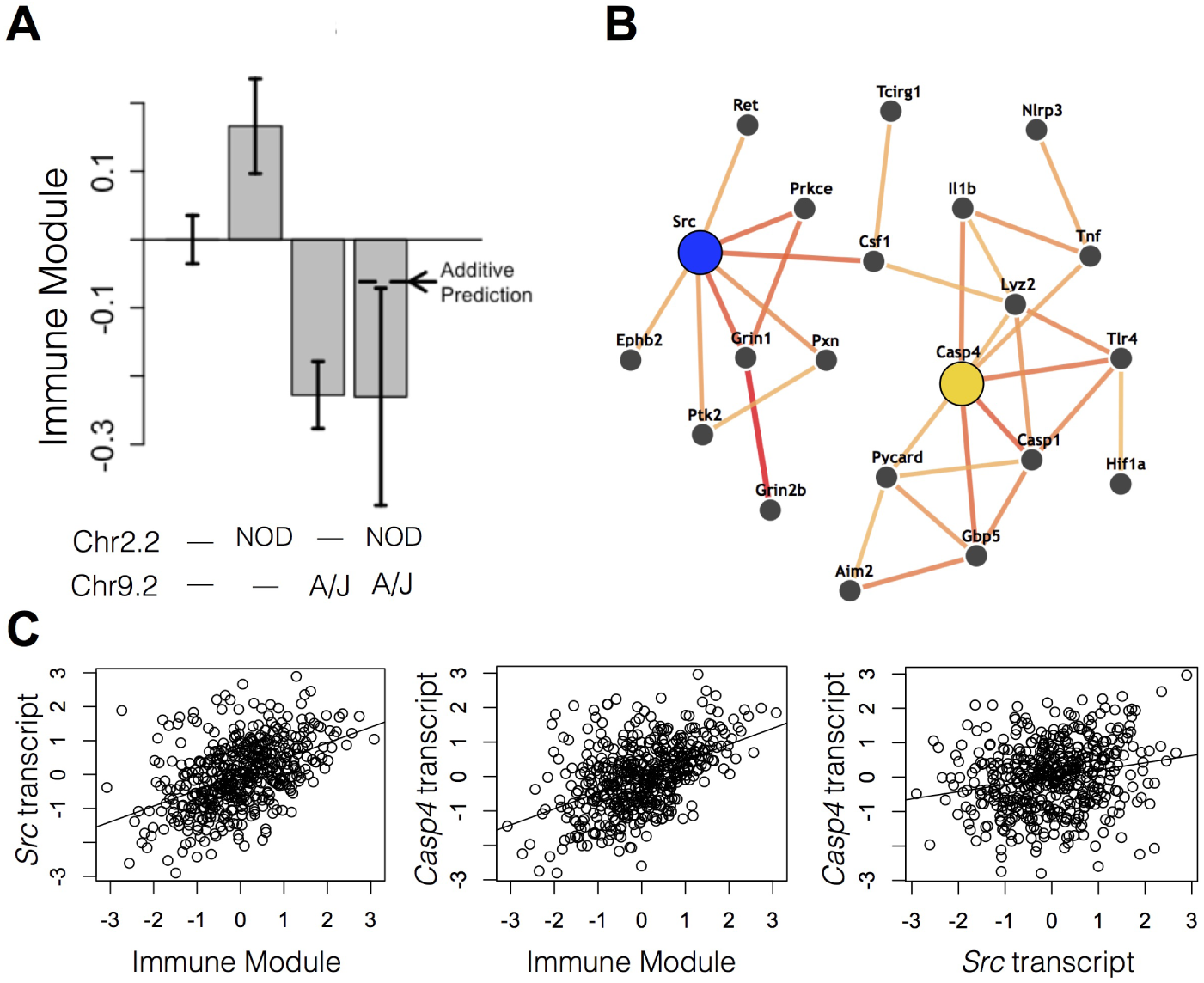
Gene prioritization for interacting QTL Chr2.2 and Chr9.2. A) The NOD haplotype at Chr2.4 and the A/J haplotype at Chr9.2 affect the immune module positively and negatively, respectively. Together, they have a negative effect similar to that of the A/J haplotype at Chr9.2. Error bars show standard error. B) Functional connections between Src and Casp4 from the IMP network. The two proteins are predicted to interact functionally by operating in related, but distinct pathways. C) The transcripts of *Src*, in Chr 2.2, and *Casp4*, in Chr 9.2, are both correlated with the immune module. The transcripts are also correlated with each other.

In addition to epistasis between genetic loci, we identified numerous QTL-sex and QTL-diet interactions. Most loci interacting with sex had effects suppressed by sex, showing greater effects in females than males. For example, the 129 allele at the Chr19.4 QTL had positive effects on triglycerides and the metabolism module. This locus possibly contains a gene that increases triglycerides through gene expression differences in metabolic pathways. Within this region there are six genes known to influence triglycerides and one of these, *Sorbs1*, had a *cis* 129-specific effect increasing *Sorbs1* expression (Figure 17A). *Sorbs1* was furthermore expressed more highly in females (*p* = 0.002) (Figure 17B), and was significantly correlated with triglyceride levels in the DO mice (*r*^2^ = 0.17, *p <* 2 × 10^-16^). Previous work has shown that mice with homozygous deletions of this gene have reduced triglyceride levels (Lesniewski *et al.* 2007). Increased expression due to the gain-of-function 129 allele is consistent with increased triglycerides in carriers, and therefore the 129 haplotype of this gene may increase risk for elevated triglyceride levels in female mice. Like sex, diet is an important factor in determining risk of metabolic disease and its related phenotypes. Diet enhanced the effects of sex suggesting that males in the DO population were more susceptible to the effects of the diet than females. This is consistent with indications that inbred male B6 mice gain more weight and have higher blood lipid profiles when fed a high-fat diet (Hwang *et al.* 2010). Multiple studies have shown interactions between genes and diet influencing factors related to traits associated with metabolic disease (for review see Ordovas (2006)). We found two QTL-diet interactions in which genetic effects on lean and/or log fat mass were suppressed in animals on the high-fat diet. Our results suggest that the effects of these variants are suppressed by the high-fat diet. Since these interactions were suppressing-coherent and therefore suggest redundancy, these QTL potentially phenocopy some of the effects of a high-fat diet.

**Figure 17.**
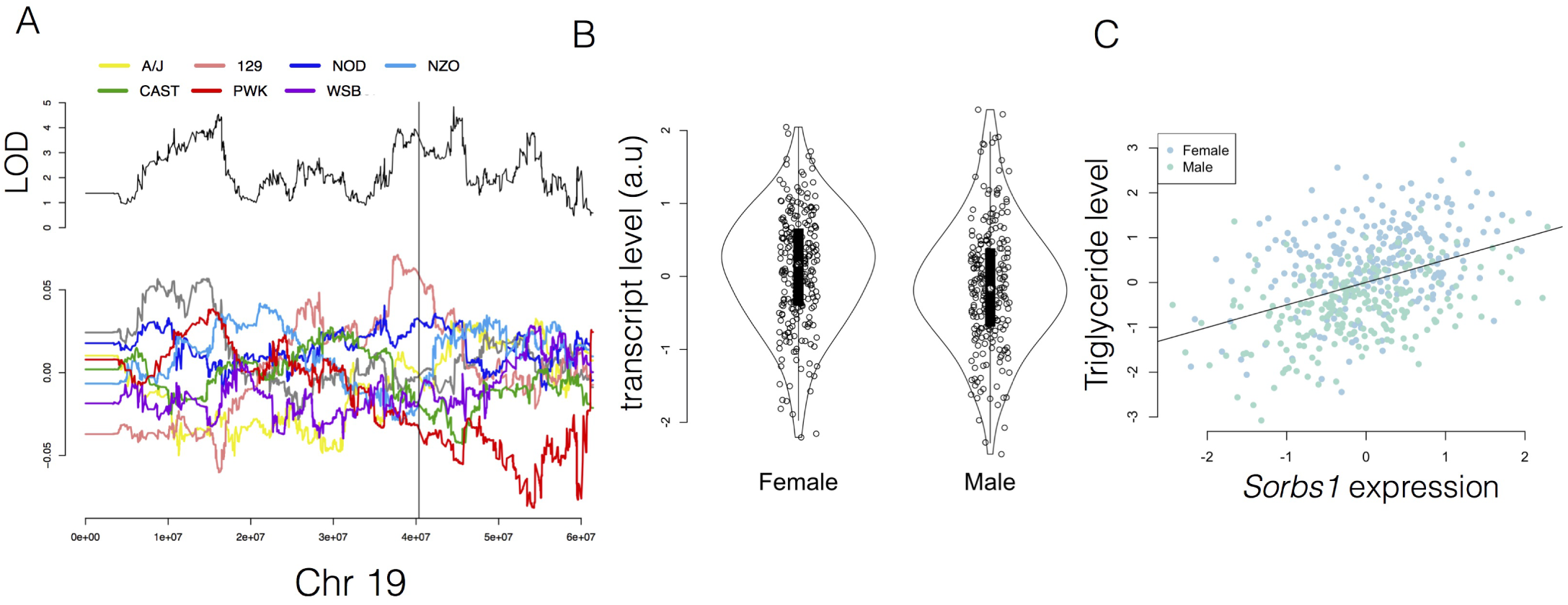
Evidence supporting a role of the 129 haplotype of Sorbs1 increasing triglyceride levels through increased transcription. A) eQTL mapping of the *Sorbs1* transcript across Chr 19. The upper panel shows LOD scores for *Sorbs1* transcript levels, with the position of the *Sorbs1* gene marked with a vertical black line. The lower panel shows haplotype effects for *Sorbs1* transcript levels. B) Transcript levels of *Sorbs1* in male and female DO mice (a.u. = arbitrary units). C) Correlation between triglyceride levels and *Sorbs1* expression (*r* = 1.7, *p <* 2 × 10^-16^). Female and male mice are shown in blue and green, respectively.

In summary, we have integrated information in physiological and transcriptional phenotypes to detect numerous genetic interactions in a relatively small DO population. Although these interactions are weak when averaged across the entire population, they can lead to large phenotypic effects in individual animals. These large phenotypic effects may be the result of incompatible recombinations between founder alleles, indicating that epistasis may be more readily detectable in multiparental populations than in traditional intercross designs between two inbred strains. By expanding the genetic diversity, multiparental populations extend the possible genetic architectures that can be studied for clinically relevant complex traits.

---

**Table S1.** Expression Modules

| Module | # Genes | Enriched GO Terms (Benjamini-adjusted p value) |
| --- | --- | --- |
| Metabolism Module | 1192 | Cellular macromolecular metabolic process ($6.3 \times 10^{-17}$)<br>Biosynthetic process ($1.6 \times 10^{-5}$) |
| Redox Module | 208 | Oxidation-reduction process ($7.7 \times 10^{-7}$)<br>Fatty acid metabolic process ($5.7 \times 10^{-2}$) |
| Immune Module | 186 | Immune system process ($5.2 \times 10^{-2}$)<br>Cell adhesion ($7.9 \times 10^{-15}$) |

**Table S2** Genotypes used in pairwise marker testing

**Figure S1.**
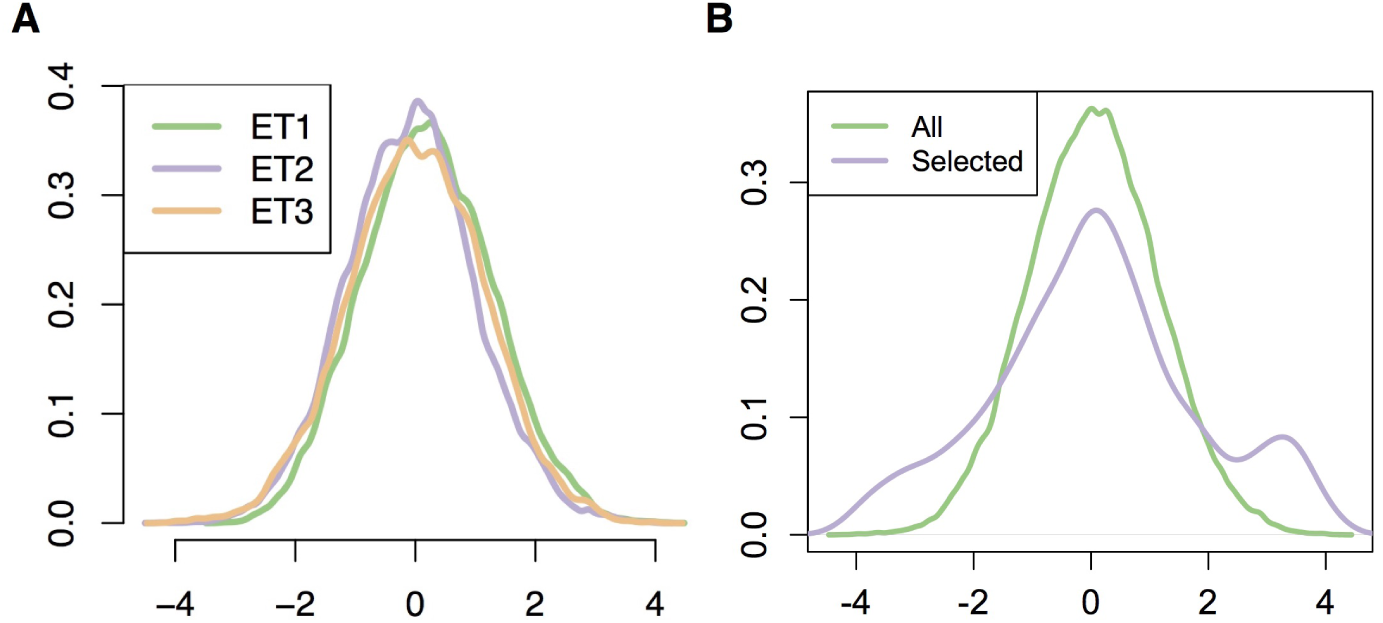
Distributions of t-statistics from allele sampling. A) Distributions of allele t-statistics across all three ETs. Distributions are comparable, indicating that no one ET was overly represented when sampling alleles with large effect sizes for pairwise testing. B) Distributions of t-statistics for all markers (green) and for the selected markers (purple) on all three ETs. The tails of the full t-statistic distribution are over-represented among the selected markers, but small t-statistics are not completely excluded.

**Figure S2.**
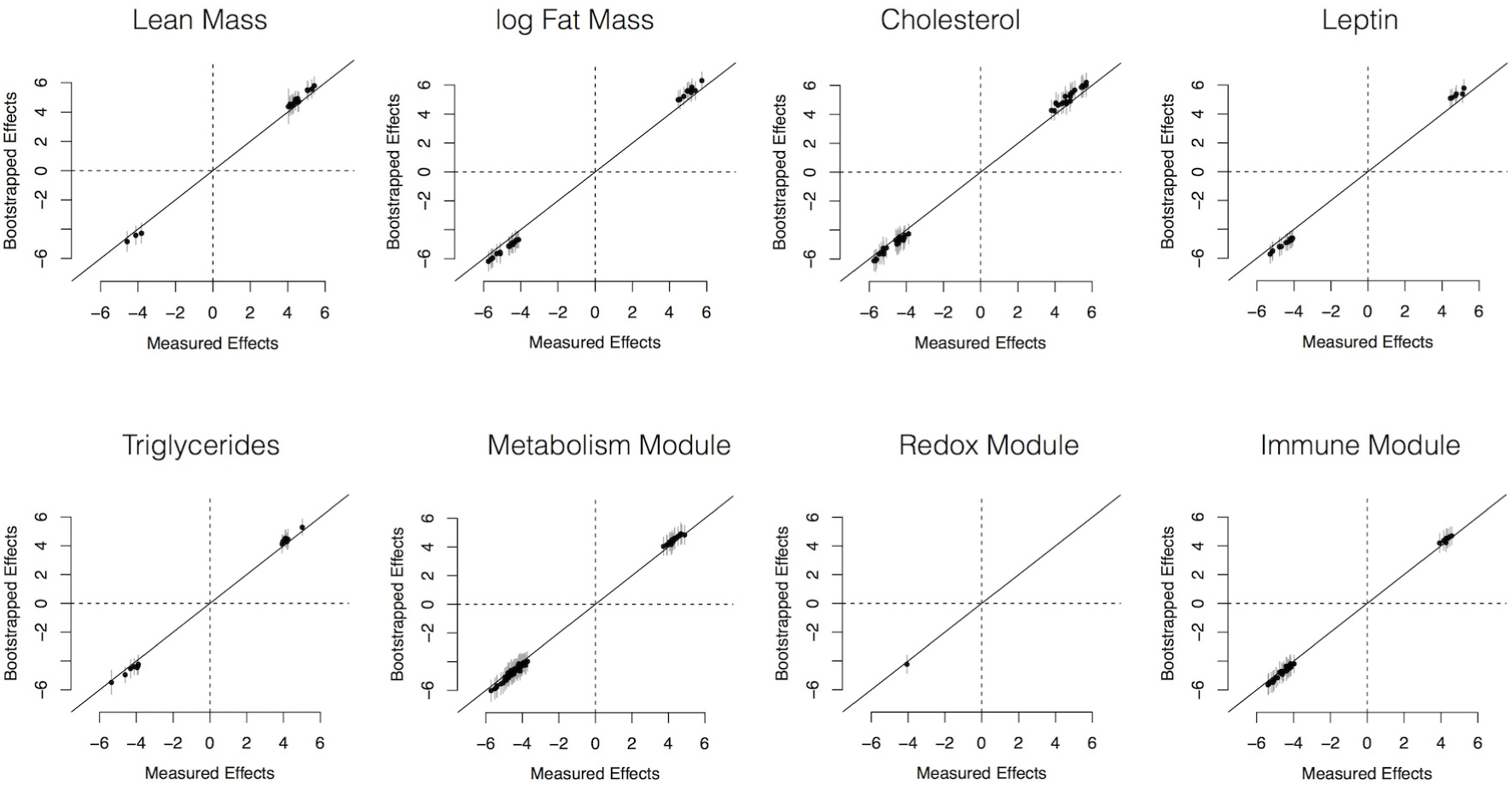
Means of bootstrapped standardized effects for all main effects in the DO network. Gray bars indicate standard deviations over 100 bootstrapped values.

**Figure S3.**
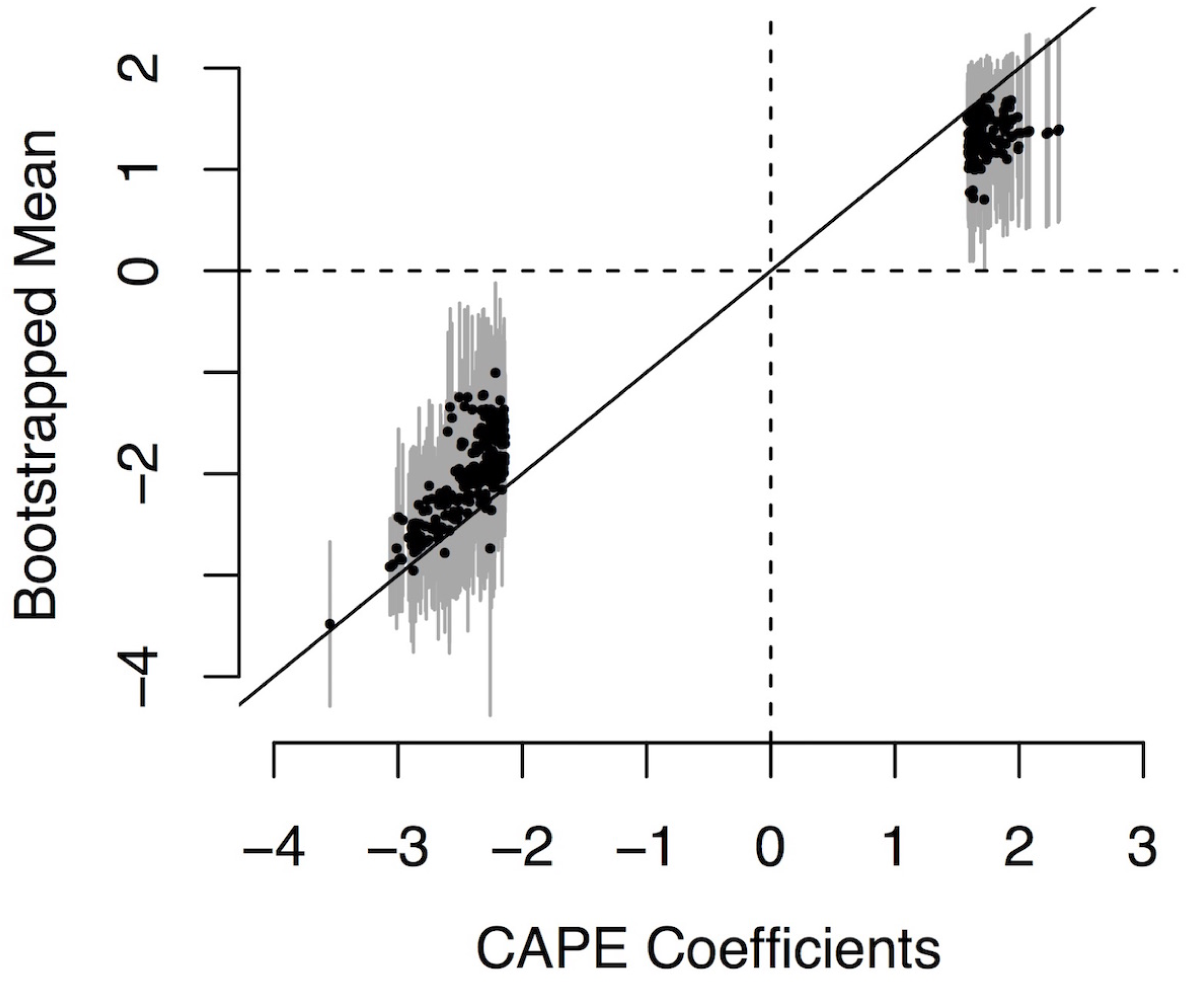
Means of bootstrapped standardized effects for all interactions in the DO network. Gray bars indicate standard deviations over 100 bootstrapped values.

